# SPARK: Bayesian Structure Learning of Enhancer-based Gene Regulatory Networks from Single-cell Multiomics Data and Prior Knowledge

**DOI:** 10.64898/2026.10.22.761021

**Authors:** Nobuaki Masaki, Shvetangi Acharya, Valeriya Malysheva, Surya Koturan, Leonardo Bottolo, Mikhail Spivakov

## Abstract

Enhancer-based gene regulatory networks (eGRNs) linking enhancers with recruited transcription factors (TFs) and downstream target genes can help decipher complex regulatory mechanisms, such as the function of GWAS variants at enhancers. Molecular assays that profile gene expression and chromatin state jointly from the same cells (such as 10x Multiome) are a rich source of input data for eGRN reconstruction. However, these assays lack mechanistic evidence for enhancer-gene and TF-enhancer links, which can be contributed by prior knowledge from other assays and analyses. Here, we present SPARK, an eGRN reconstruction framework that formally combines single-cell multiome data with mechanistic prior knowledge. SPARK represents eGRNs as directed acyclic graphs (DAGs) and learns them using Bayesian inference, with a prior on the network structure constructed from heterogeneous sources of evidence. In a simulation study, we show that SPARK benefits from prior knowledge and outperforms existing DAG-based and regression-based algorithms. Finally, we use SPARK to study the regulatory circuitry of enhancers harboring immune disease GWAS variants in human primary monocytes, revealing known and novel regulatory links as candidates for experimental validation.

## 1 Introduction

DNA cis-regulatory elements such as enhancers orchestrate cell state transitions, particularly in development and in response to environmental signals. Enhancers recruit combinations of sequence-specific transcription factors (TFs) that regulate their chromatin accessibility and activity. The activating and potentially repressive signals from enhancers modulate the transcriptional state of their target genes located up to millions of base pairs away through chromosomal interactions that bring these elements into physical proximity in the 3D space of the nucleus [1, 2].

Enhancer-based gene regulatory networks (eGRNs) formalize this regulatory logic as a graph, in which enhancers, the TFs recruited to them, and their downstream target genes are represented as nodes, and the regulatory relationships between them as edges, enabling their formal reconstruction and interrogation from multiomics data [3–7]. In particular, the explicit representation of enhancers as nodes makes eGRNs attractive tools for interpreting the functional effects of GWAS variants that are highly enriched within these elements [4, 8].

Modern multiomics technologies, such as 10x Multiome, which jointly profile chromatin accessibility and gene expression across single cells, provide a rich source of input data for eGRN reconstruction, as correlations between these properties at specific genes and enhancers can provide statistical evidence for their regulatory relationships. However, only a subset of correlations in the data represent direct regulatory relationships, with the rest reflecting spurious correlations or indirect effects. The high sparsity and technical noise inherent in single-cell data can also weaken detectable associations even when direct regulatory relationships exist.

Incorporating mechanistic evidence for regulatory relationships into eGRN inference can help address these challenges [9]. Two complementary sources of prior information are particularly relevant for this purpose: TF binding information from sequence-based predictions or biochemical assays such as ChIP-seq or CUT&RUN, and enhancer-promoter contact data obtained from chromosome conformation capture assays. Multiple computational tools have combined multiomics data with one or more prior knowledge sources for eGRN reconstruction, often using them to constrain the set of candidate regulatory interactions in approaches ranging from correlation analysis (scMEGA) [5] to linear and non-linear regression (Pando, SCENIC+) [3, 6], graphical LASSO (SCEG-HiC) [10], and deep learning (LINGER) [7]. However, although cis-regulatory relationships are inherently directional, these methods typically capture statistical associations or predictive relationships rather than directed regulatory effects. This limits the ability to identify true regulatory hierarchies, prioritize functional regulatory elements, and accurately simulate the consequences of specific perturbations.

Directed acyclic graphs (DAGs) provide a natural framework for representing directed effects between TFs, enhancers, and target genes in eGRNs, while imposing a coherent acyclic structure that enables their intuitive interpretation and formal analysis. However, the formal inclusion of regulatory information as a prior on the network structure remains an open problem in DAG-based eGRN inference.

Here, we present SPARK, a Bayesian DAG inference framework for learning eGRNs from observational single-cell multiome data. SPARK constructs a prior on the network structure from heterogeneous sources of evidence and uses it to provide mechanistic support to candidate regulatory interactions. We validate our approach on simulated single-cell multiome data. As a use example, we apply SPARK to predict the regulatory circuitries of enhancers harboring immune disease risk variants in human primary monocytes, revealing known and potentially novel regulatory relationships between these enhancers, their upstream TF regulators, and downstream target genes.

## 2 Results

### 2.1 Summary of SPARK

SPARK (Structure Learning with Priors Augmented by Regulatory Knowledge) is a Bayesian framework that reconstructs enhancer-based gene regulatory networks (eGRNs) from observational singlecell multiome data, in which chromatin accessibility (scATAC-seq) and gene expression (scRNA-seq) are profiled in the same cell. SPARK takes two inputs: (i) a matrix of counts these assays produce for *m* features (genes and enhancers) across *N* cells or metacells (Figure 1A), and (ii) a topological prior that labels plausible regulatory relationships by the sources of evidence supporting them (Figure 1B).

**Figure 1.**
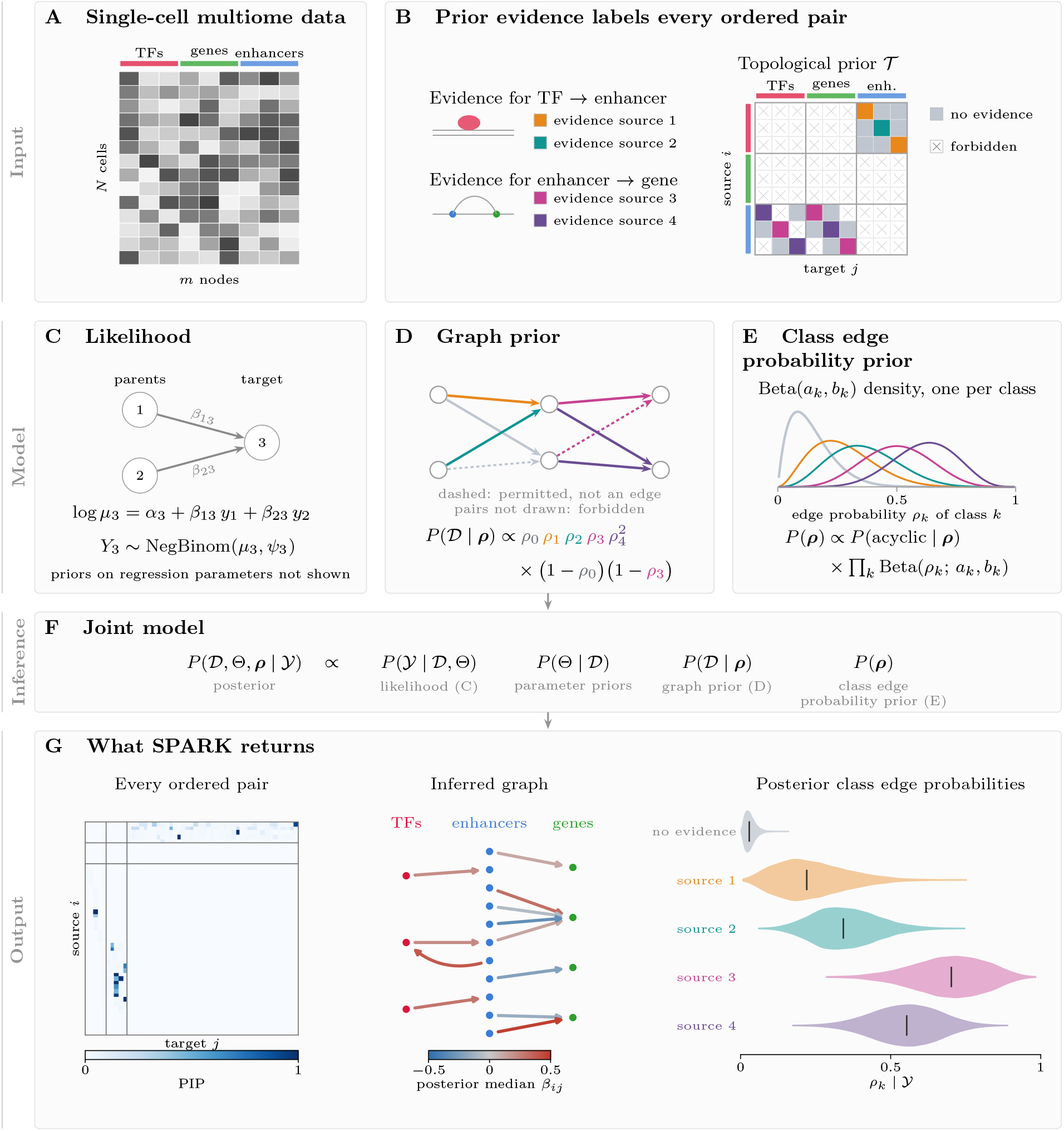
Overview of SPARK. **A.** Single-cell multiome data. Counts of chromatin accessibility and gene expression over *m* nodes and *N* cells. **B**. Topological prior. Each ordered pair of nodes is labeled by the evidence source supporting it. Permitted pairs supported by no source form the no-evidence class, and forbidden pairs never appear as edges in the inferred DAG. **C**. Likelihood. Each node follows a negative binomial regression on the counts of its parents. **D**. Graph prior. Each permitted pair contributes a factor of *ρ*_*k*_ if it is an edge and 1 −*ρ*_*k*_ if not, where *ρ*_*k*_ is the edge probability of its class. The equation shows this product for the DAG drawn. **E**. Prior on the class edge probabilities. The user specifies a Beta density for each *ρ*_*k*_, and their product is adjusted for acyclicity. **F**. Joint posterior of the DAG, regression parameters, and class edge probabilities, sampled by MCMC. **G**. Output. The posterior inclusion probability (PIP) of each edge, the fraction of sampled DAGs containing it, and the posterior of each class edge probability. A network can be drawn by keeping the edges whose PIP exceeds a threshold, coloring them by their posterior median direct effect *β*_*ij*_ representing the magnitude and direction of the effect.

SPARK represents eGRNs as directed acyclic graphs (DAGs) whose nodes can be enhancers, their upstream TF regulators, and their downstream target genes, with directed edges between them representing regulatory effects. To quantify the sign and magnitude of each regulatory effect, SPARK’s likelihood models the read count of each node using a negative binomial regression on the counts of its parental nodes in the graph (Figure 1C).

SPARK builds the topological prior from knowledge about plausible TF-enhancer and enhancergene links, typically obtained from different sources of evidence (here, we used TF ChIP-seq data and in silico predictions for TF-enhancer links and linear distance and Capture Hi-C data for enhancer-gene links). Based on this information, SPARK labels each plausible regulator-target pair according to the source of evidence, if any, that supports their relationship. The pairs supported by each source form an evidence class, and the plausible pairs that no source supports form the no-evidence class. In addition, we assume that some relationships are mechanistically implausible, including genes directly regulating the expression of other genes and enhancers directly regulating genes on a different chromosome or too far away from them on the same chromosome. These relationships are labeled as forbidden and will never appear as edges in the final eGRN (Figure 1B).

SPARK’s graph prior expresses how plausible a network is before considering the observed data, based on the evidence available for each potential edge. In particular, it assigns a high probability to networks that include pairs supported by reliable evidence sources and exclude pairs for which there is little or no supporting evidence. The reliability of each evidence source is captured by the class edge probabilities, which govern how likely a pair supported by each source is to be included as an edge in the resulting network (Figure 1D). The prior on the class edge probabilities encodes the user’s belief in the reliability of each source (Figure 1E).

The Markov chain Monte Carlo (MCMC) algorithm in SPARK samples DAGs, the parameters of the negative binomial regression used in the likelihood, and the class edge probabilities jointly from their posterior distribution (Figure 1F). The resulting posterior samples can be used to infer which regulatory effects are present, the magnitude and direction of effect of each putative regulator on its downstream nodes, as well as the informativeness of each prior source of evidence (Figure 1G).

### 2.2 Using topological prior information improves DAG reconstruction

We first tested SPARK on data simulated from known regulatory networks, assuming the topological prior as the source of ground-truth DAGs. To this end, we used the topological prior assembled for human M-CSF-stimulated monocytes. We focused on a subset of this prior comprised of 51 features located on chr2 (5 TFs, 41 enhancers, and their 5 plausible target genes). In total, this subset contained 302 plausible regulatory interactions, 21 of which were supported by JASPAR2024 motif matches [11] and 13 by Promoter Capture Hi-C (PCHi-C) contacts [12]; the remaining plausible interactions were assigned to the no-evidence class (Figure 2A). From these 302 permitted pairs, we generated ten ground-truth DAGs, whereby the inclusion probability of each pair as an edge in the DAG depended on its source of evidence. We defined these probabilities as the means of the expert-elicited Beta densities for each source, 0.286 for JASPAR2024 and 0.606 for PCHi-C; we assumed an edge probability of 0.05 for the remaining plausible pairs (Table S1). We then sampled the direct effect *β*_*ij*_ of each included edge, corresponding to the sign and magnitude of the regulatory effect, from a normal distribution. From each of the ten DAGs, we then simulated a scMultiome dataset of *N* = 50 cells, whereby the counts of each feature (gene or enhancer) depended on those of their direct regulators according to the DAG. To achieve this, we drew the counts of each feature from a negative binomial distribution whose mean depended on the counts of its parents in the DAG, weighted by the direct effects along the edges connecting them (Methods).

**Figure 2.**
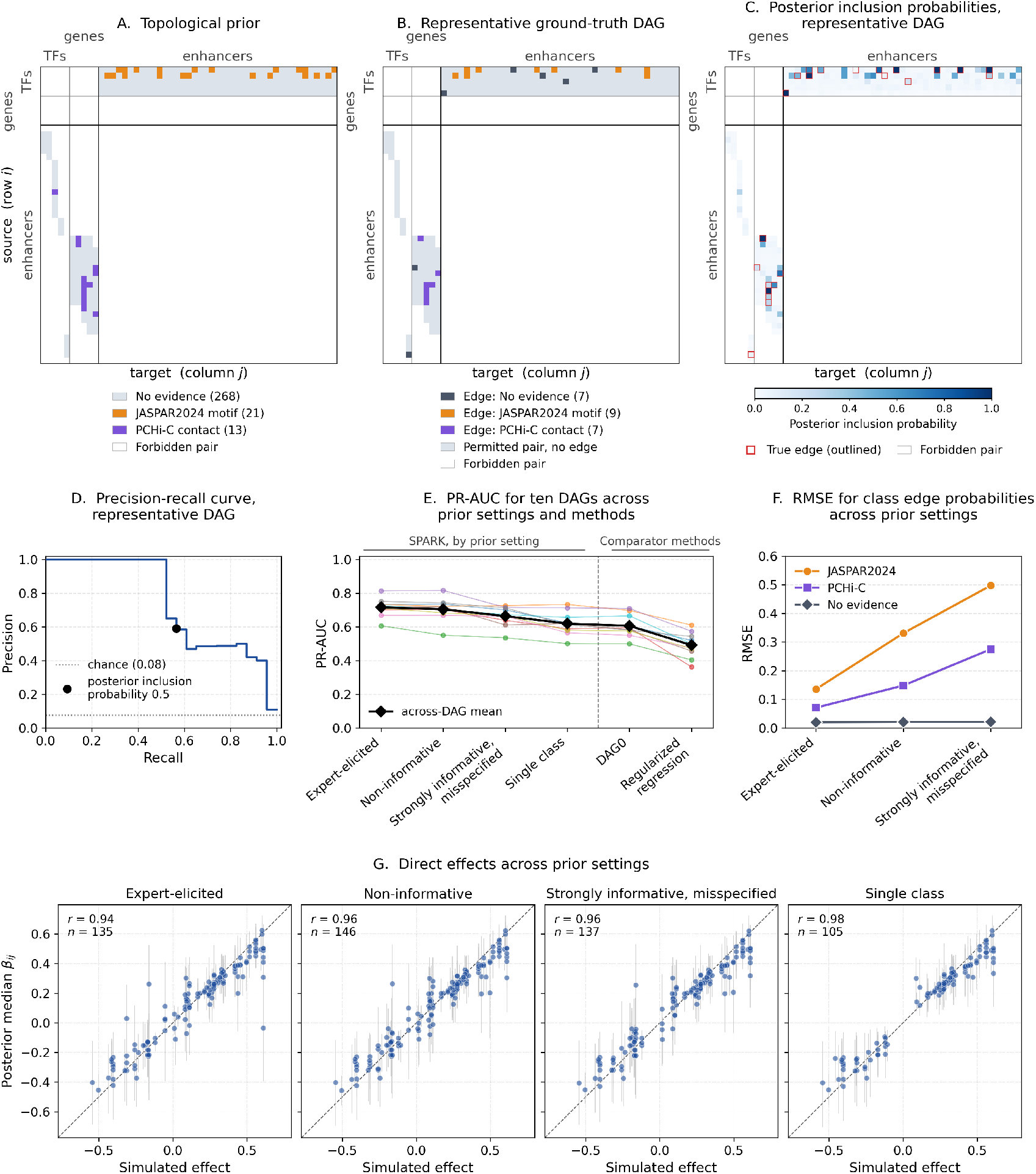
Edge recovery on simulated data. **A.** Topological prior for M-CSF-stimulated human monocytes, restricted to the 51 TFs, enhancers, and plausible target genes on chromosome 2. Each box is an ordered pair of nodes. **B**. Representative ground-truth DAG (DAG 7), one of ten DAGs drawn from the topological prior in A. **C**. Posterior inclusion probabilities (PIPs) inferred by SPARK under the non-informative prior setting from 50 cells simulated from the representative DAG. **D**. Precision-recall curve for the representative DAG. The dotted line is the prevalence of true edges among the permitted pairs, and the black point is the precision and recall at a PIP threshold of 0.5. **E**. PR-AUC of SPARK under each prior setting, DAG0, and the regularized regression from CellOracle. Each colored line is one of the ten ground-truth DAGs. **F**. RMSE of the posterior median class edge probabilities against their generating values over the ten DAGs, under the three prior settings with evidence classes. **G**. Posterior median of each direct effect *β*_*ij*_ against its simulated value under each prior setting, for the true edges with a PIP of at least 0.5, pooled across the ten DAGs. Gray lines are 95% central credible intervals. Each plot reports the number of these edges *n* and the Pearson correlation *r* between the posterior medians and the generating values.

We ran SPARK on each of the ten simulated datasets under four prior settings (Table 1). In the rest of this section, we examine the result of SPARK network inference under the non-informative prior setting on one of the ten datasets, comparing the resulting eGRN to the ground-truth DAG from which the dataset was simulated. We then extend the analysis to all ten datasets and the other three prior settings to assess how these settings affect the accuracy of network reconstruction. Next, we benchmark SPARK against a Bayesian DAG reconstruction method for count data (DAG0 [13]) and Bayesian ridge regression used in CellOracle [14]. Finally, we assess how accurately SPARK estimated parameters used to generate the ground-truth DAGs and simulated datasets.

**Table 1.** Prior settings in the simulation study. Each setting specifies the evidence classes given to SPARK and the Beta prior on the edge probability of each class. The last row gives the edge probabilities used to generate the ground-truth DAGs.

| Setting | Evidence classes | Prior on $\rho_1$<br>(JASPAR2024) | Prior on $\rho_2$<br>(PCHi-C) | Prior on $\rho_0$<br>(no evidence) |
| --- | --- | --- | --- | --- |
| (1) Non-informative | JASPAR2024,<br>PCHi-C | Beta(1, 1) | Beta(1, 1) | Beta(1, 1) |
| (2) Expert-elicited | JASPAR2024,<br>PCHi-C | Beta(2.20, 5.50),<br>mean 0.286 | Beta(5.68, 3.70),<br>mean 0.606 | Beta(1, 1) |
| (3) Strongly<br>informative,<br>misspecified | JASPAR2024,<br>PCHi-C | Beta(16.8, 4.2),<br>mean 0.80 | Beta(3.25, 9.75),<br>mean 0.25 | Beta(1, 1) |
| (4) Single class | None (all 302<br>permitted pairs in<br>the no-evidence<br>class) | — | — | Beta(1, 1) |
| Generating edge<br>probability |  | 0.286 | 0.606 | 0.05 |

We first focused on the results of SPARK inference from data simulated from DAG 7, a representative ground-truth DAG with 23 edges (Figure 2B; Table S2), using non-informative priors on the edge probability of each class. We compared the resulting eGRN (Figure 2C) at a PIP threshold of 0.5 with the edges in DAG 7. In this setting, SPARK achieved a precision of 0.59 (13 correct out of 22 called edges) and a recall of 0.57 (13 of the 23 true edges recovered). The area under the precision-recall curve (PR-AUC), calculated across all PIP thresholds, was 0.742 (Figure 2D; Table S2). Expanding the analysis to data simulated from the other nine DAGs, we found that, on average, they achieved a similar performance (mean PR-AUC = 0.704 *±* 0.021 SE for the non-informative prior setting, Figure 2E; Table S2).

We next explored how prior settings other than non-informative priors on the edge probability for each class influenced this result. For this, we considered three additional scenarios (Table 1): (2) expert-elicited priors, whose Beta densities had means equal to the generating edge probabilities; (3) strongly informative but misspecified priors, whose Beta densities had means of 0.80 for the JASPAR2024-supported class and 0.25 for the PCHi-C-supported class; and (4) single class, with all 302 permitted pairs assigned to the no-evidence class. Averaged over the ten ground-truth DAGs, SPARK’s PR-AUC was the highest under the expert-elicited setting (0.716), followed by the non-informative (0.704), strongly informative but misspecified (0.664), and single-class (0.619) settings (Figure 2E).

We used these simulated datasets to benchmark SPARK against two other methods: a Bayesian DAG reconstruction method for count data that does not take topological prior knowledge (DAG0), and Bayesian ridge regression used in a popular GRN reconstruction tool, CellOracle (Methods). We found that the mean PR-AUCs of DAG0 (0.606) and CellOracle-style Bayesian ridge regression (0.492) were lower than SPARK’s under every prior setting (Figure 2E).

We next assessed how well SPARK estimated the informativeness of each class at different prior settings. For this, we compared posterior median edge probabilities for each class with the probabilities at which pairs in this class were included as edges when simulating the ground-truth DAGs. The expert-elicited prior setting showed the best performance, followed by the non-informative and the strongly informative but misspecified prior settings (Figure 2F; Table S3).

Finally, we assessed how well SPARK estimated the direct effects, which give the sign and strength of each regulatory effect. For this comparison, we retrieved the posterior median direct effects of the ground-truth edges with a PIP of at least 0.5, pooled across the ten DAGs (105 to 146 of the 252 edges, depending on the prior setting), computing each median over the posterior samples in which the edge was present (Methods). Under all four prior settings, these posterior medians were strongly correlated with their corresponding generating values used to simulate the scMultiome data (*r* = 0.94 to 0.98; Figure 2G). The model also accurately estimated the other two parameters used in data generation: node intercepts *α*_*j*_ and overdispersions *ψ*_*j*_ (Figure S1).

In conclusion, integrating topological prior information improves SPARK’s edge recovery, even when prior beliefs on the informativeness of each source are misspecified. SPARK outperformed DAG0 and CellOracle-style Bayesian ridge regression in the accuracy of network reconstruction under every prior setting and accurately estimated the sign and strength of each discovered regulatory effect.

### 2.3 SPARK links enhancers harboring immune disease GWAS variants to their regulators and target genes in M-CSF-stimulated monocytes

As an illustration of SPARK analysis on real data, we used it to study the regulatory circuitry of enhancers harboring immune disease GWAS variants (“immGWAS enhancers”) in human primary monocytes stimulated with monocyte colony-stimulating factor (M-CSF) for 16h. We generated 10x Multiome profiles of 3141 M-CSF-stimulated CD14^+^ monocytes, capturing single-cell gene expression and chromatin accessibility in the same cells, and aggregated them into 83 metacells to mitigate data sparsity (Methods).

We centered the analysis on a set of input features comprising 14 immGWAS enhancers, selected based on their overlap with fine-mapped immune GWAS loci from the Open Targets Platform [15]. As candidate target genes of these enhancers, we considered 17 genes located in their 1 Mb proximity. We further included 57 TFs as putative upstream regulators (2/57 TFs were also candidate target genes of immGWAS enhancers) and 291 additional enhancers located within 1 Mb of the TSS of any selected gene. In total, we obtained 377 putative network nodes that were used as input for SPARK (Methods).

We next assembled the topological prior for the selected nodes. As sources of prior evidence for TF-enhancer edges, we used monocyte and macrophage TF ChIP data from ChIP-Atlas [16] (207 links) and JASPAR2024 motif matches (1,747 links). As evidence for enhancer-gene pairs, we used linear proximity (enhancer within the gene body, 17 links) and contacts detected in PCHi-C in monocytes and macrophages [12] (51 links). We allowed a further 15,785 edges linking each TF to each enhancer and each enhancer to any proximal gene (within 1 Mb from the TSS), which were not supported by prior evidence, placing them in the no-evidence class. We prohibited the remaining 123,945 of the 141,752 combinatorially possible edges between the 377 selected nodes. We ran SPARK under the expert-elicited prior setting with 20 parallel-tempered chains for 5,250,000 iterations, until the traces of the log-posterior and of the number of edges in the sampled DAGs had flattened, and discarded the first 3,000,000 iterations as burn-in (Methods). These traces, together with the acceptance rates of the sampler’s updates and of the swaps between chains, are shown in Figures S2 to S5.

Figure 3A shows the inferred network, at a PIP threshold of at least 0.5 used throughout the analysis as evidence of regulatory relationships. AFF3, NFIA, and PPARG emerged as the top 3 regulators in the network, with 133, 119, and 87 outgoing edges, respectively, jointly comprising just over 60% of the TF-enhancer edges in the network. Notably, these edges varied widely in their degree of prior support: from 35.3% of edges originating from NFIA to 0% for those originating from AFF3, for which no prior evidence of binding was available. Overall, while only 2,022 of the 17,807 permitted network edges (11%) were supported by prior evidence, they accounted for 134/605 (22%) of predicted edges, suggesting that prior evidence was generally informative for the placement of edges in the reconstructed eGRN. Specifically, both prior sources of enhancer-gene links were highly informative (linear proximity: 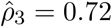, 95% CrI: [0.45, 0.94]; PCHi-C: 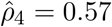, 95% CrI: [0.40, 0.75]), while JASPAR2024 as a prior for TF-enhancer links was less informative 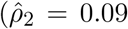, 95% CrI: [0.07, 0.12]), likely reflective of imperfections in TF binding inference from sequence. Finally, ChIP-Atlas was as informative (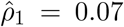, 95% CrI: [0.03, 0.12]) as the no-evidence class (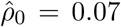, 95% CrI: [0.06, 0.07]). This is likely due to the small number of TFs (EGR1, EGR2, and PPARG) included in this data source. Furthermore, EGR1, accounting for the majority of pairs in ChIP-Atlas (180*/*207), gave rise to only four prior-supported edges in the final eGRN, and was therefore likely inconsistent with the observed scMultiome data.

**Figure 3.**
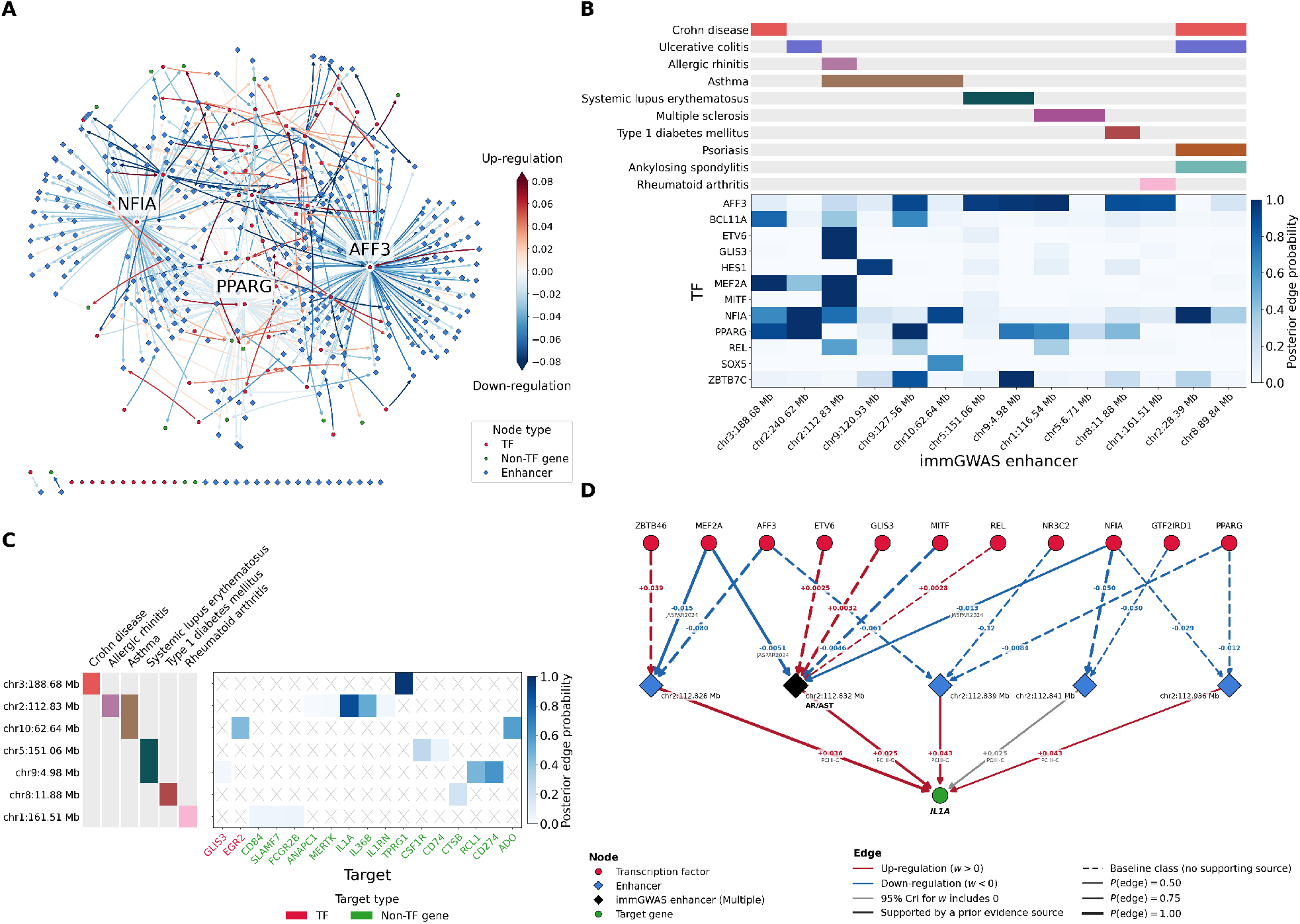
The inferred immGWAS enhancer-centered network for M-CSF stimulated monocytes. **A.** The inferred network, obtained by drawing every edge whose posterior inclusion probability (PIP) is at least 0.5. Edge color gives the posterior median direct effect. The three TFs with the most edges are labeled. **B**. PIPs of the edges from TFs (rows) to the 14 immGWAS enhancers (columns), for the 12 TFs with a PIP of at least 0.5 to at least one of these enhancers. Strips above the heatmap mark the disease categories of each enhancer’s GWAS variant. **C**. PIPs of the edges from immGWAS enhancers (rows) to their candidate target genes (columns), for the 7 immGWAS enhancers with at least one target permitted by the topological prior. Strips on the left mark the disease categories of each enhancer’s GWAS variant. Crossed cells are pairs the topological prior forbids. **D**. The cis-regulatory unit of *IL1A*, obtained by drawing every edge into *IL1A* whose PIP is at least 0.5, and then every such edge into the enhancers regulating *IL1A*. The black enhancer is the immGWAS enhancer `chr2:112.83 Mb`;, which carries a variant from an allergic rhinitis and asthma study (AR/AST). Edge labels give the posterior median direct effect and, for prior-supported edges, the supporting evidence source. Edges are colored red or blue only when the 95% credible interval of the direct effect excludes 0.

We next considered the sign of the inferred direct effects, reflecting either positive or negative regulatory effects. As expected, the majority of enhancer-to-gene effects were positive (36*/*52, 69%). In contrast, three-quarters of TF-to-enhancer effects were negative (415*/*553, 75%); however, this varied significantly by TF (Table S4). For example, 80% of PPARG’s effects on enhancers were negative, potentially corresponding to its activity in the ligand-unbound state [17], while 75% of MEF2A effects were positive, consistent with its well-characterized activator function [18]. Notably, the proportion of negative effects was similar for the edges supported and unsupported by prior knowledge (74.5% versus 75.2%, respectively).

Overall, 12/57 analyzed TFs were predicted to regulate the 14 immGWAS enhancers (Figure 3B). Of these, AFF3 was predicted to regulate 6/14, NFIA and PPARG 5 each, with the remaining TFs regulating no more than 2 immGWAS enhancers. Enhancers regulated by AFF3 carried variants for systemic lupus erythematosus, multiple sclerosis, type 1 diabetes, rheumatoid arthritis, and asthma, consistent with *AFF3* ‘s localization within a susceptibility locus for rheumatoid arthritis and type 1 diabetes [19, 20]. Enhancers regulated by NFIA harbored variants for Crohn’s disease, ulcerative colitis, and asthma, consistent with *NFIA*’s known association with the combined asthma and allergic rhinitis phenotype [21]. Finally, enhancers regulated by PPARG carried variants for Crohn’s disease, ulcerative colitis, asthma, systemic lupus erythematosus, and multiple sclerosis. This is consistent with PPARG’s function as an anti-inflammatory nuclear receptor, whose expression is reduced in the colon of ulcerative colitis patients [22, 23], as well as the emerging potential of PPARG agonists in the therapy of this disease [23].

At a PIP threshold of 0.5, SPARK predicted five connections from immGWAS enhancers to their putative target genes, including the inflammatory cytokine genes *IL1A* and *IL36B*, the immune checkpoint ligand gene *CD274* (*PD-L1* ), as well as the *ADO* and *TPRG1* genes whose function in the immune system is less well understood (Figure 3C). *IL1A*, regulated by an enhancer harboring an allergic rhinitis/asthma (AR/AST) risk variant in the eGRN, has a known role in asthma and COPD pathogenesis [24]. The same immGWAS enhancer was also connected to *IL36B*, which encodes IL-36*β*, a cytokine elevated in the sputum of patients with neutrophilic asthma [25]. Finally, *CD274*, regulated by an enhancer harboring a systemic lupus erythematosus risk variant, was previously shown to have reduced expression in monocytes in lupus patients [26].

In addition to an immGWAS enhancer, *IL1A* was predicted to be regulated by four others, the most of any gene in the eGRN, with all five supported by PCHi-C contacts and were inferred to be regulated by 11 TFs through 16 edges (Figure 3D). However, only 3 of these 16 edges are supported by a prior evidence source, all by JASPAR2024 motif matches, pointing to either imperfect binding prediction or potential indirect effects. Notably, some of both JASPAR2024-supported and unsupported regulatory relationships are consistent with prior literature. For example, PPARG binds near *IL1A* in ChIP-seq data compiled by GTRD [27], and its activation inhibits the production of IL-1 and other inflammatory cytokines in monocytes [28], in line with the negative direction of PPARG-to-*IL1A* relationship in the eGRN. PPARG, REL, MEF2A, and ETV6 were also predicted to regulate *IL1A* in the tissue-specific regulatory networks of Marbach *et al*. [29]. However, for the remaining seven TFs, including NFIA, whose binding to the *IL1A*-connected immGWAS enhancer was supported by a JASPAR2024 motif match, we found no previously reported links. Finally, we compared the upstream regulatory networks between *IL1A* (Figure 3D) and its antagonist *IL1RN* (Figure S6). We found that NFIA had JASPAR2024-supported links to the enhancers of both genes, but with opposite directions of effect, consistent with their opposing biological functions.

In conclusion, the analysis of immGWAS enhancer-centered directed eGRN in M-CSF-stimulated human primary monocytes has revealed known, plausible, and unexpected regulatory relationships, demonstrating the potential of our SPARK framework to generate testable hypotheses for further experimental validation.

## 3 Discussion

Here we presented SPARK, a Bayesian DAG inference framework that reconstructs eGRNs by combining statistical evidence from observational single-cell multiome data with mechanistic evidence from heterogeneous sources, which it encodes as a prior on the network structure. In a simulation study, SPARK consistently outperformed DAG0 [13] and CellOracle-style Bayesian ridge regression [14], particularly when evidence sources underlying the network structure were provided. Applied to single-cell multiome data from M-CSF-stimulated monocytes, SPARK recovered known, plausible, and unexpected regulatory relationships around immGWAS enhancers, generating testable hypotheses for experimental validation.

A key motivation for SPARK was to infer directed regulatory effects, which are informative for identifying regulatory hierarchies and predicting the effects of perturbing individual TFs or enhancers. Accurately recovering these effects from observational data relies on the identifiability of SPARK’s negative binomial DAG model, which requires, among other assumptions, that no unobserved features regulate two or more of the nodes in the DAG [13]. This assumption rarely holds in practice, but we attempted to reduce latent confounding by including likely regulators as nodes: in our analysis of single-cell multiome data from M-CSF-stimulated monocytes, we included all TFs with highly variable expression and all highly accessible enhancers located in the proximity of the selected genes, resulting in a network of 377 nodes.

The number of possible DAGs grows super-exponentially with the number of nodes, making reconstruction of large networks computationally challenging. The topological prior keeps SPARK network inference tractable for networks with hundreds of nodes by prohibiting mechanistically implausible regulatory relationships, which restricts the space of DAGs SPARK’s MCMC algorithm must explore.

Many existing methods for eGRN or GRN inference go further and restrict the network to pairs supported by prior evidence. For example, SCENIC+ links a TF only to regions enriched for its motif [3] and CellOracle allows a TF to regulate a gene only if the TF’s motif is found in the gene’s promoter or linked enhancers [14]. However, this can produce spurious associations between pairs that are supported by prior evidence but are correlated only because they share a regulator whose effects on the pair are excluded by these constraints. Permitting non-prior-supported TF-enhancer edges in SPARK’s network inference helps mitigate these spurious associations. However, as predicted TF-enhancer edges without prior support may reflect both indirect effects and gaps in prior knowledge about direct relationships, they may need to be considered separately in downstream analysis.

In our use example, we “read” dependency relationships directly from the inferred graph. However, SPARK’s DAG eGRNs are well suited to formally predicting the effects of perturbations using do-calculus [30]. This approach makes it possible to express perturbation effects as the fold changes in the mean expression or accessibility of each node downstream of the perturbed node relative to its unperturbed state. As a Bayesian method, SPARK can propagate its posterior uncertainty about the network structure and regression parameters into these predicted effects. Implementing this framework is the subject of future developments.

Because SPARK uses DAGs to represent eGRNs, it cannot capture feedback loops, which are common in gene regulation. Methods based on linear stochastic differential equations, such as Dictys [31], allow feedback loops by treating single-cell expression as draws from the steady state of a stochastic process. However, it was shown that in continuous Lyapunov models, which Dictys is based on, the directed graph cannot in general be identified from observational data [32]. Dictys partly addresses this by restricting the graph to candidate edges supported by TF footprints. SPARK, by contrast, fits a negative binomial DAG model whose structure is fully identifiable from observational data under certain conditions [13].

In this study, we restricted the sources of prior evidence to close genomic proximity and PCHi-C contacts for enhancer-gene edges, and to motif matches and TF ChIP data for TF-enhancer edges. Our results suggest that adding further informative sources of prior evidence should improve SPARK’s ability to recover true regulatory relationships. For example, TF binding predictions from deep learning models trained on sequence data such as BPNet [33], Enformer [34], and AlphaGenome [35], as well as enhancer-promoter pairs inferred from bulk data using approaches such as ABC [36] and ENCODE-rE2G [37] and from single-cell data using methods such as CICERO [38] and sc-E2G [39] can be leveraged as additional sources of prior evidence. SPARK’s ability to learn how informative each evidence source is for identifying regulatory effects makes it particularly well suited to evaluating the applicability of these predictions in eGRN reconstruction. Finally, as CRISPR perturbation screens with single-cell multiome readouts are becoming feasible [40–44], extending SPARK to learn jointly from perturbational and observational datasets is a promising further direction to improve the accuracy of reconstructed eGRNs and the directed effects inferred in them.

## 4 Methods

### 4.1 DAG formulation, likelihood, and priors on the regression parameters

Let *D* = (*V, E*) denote a DAG, where *V* = *{*1, 2, …, *m}* is the set of nodes and *E* ⊆ *{*(*i, j*) ∈ *V × V* : *i* ≠ *j}* the set of directed edges. We say *u* is a parent of *v* in *D* = (*V, E*) if there is an edge (*u, v*) ∈ *E*, and denote the parent set of *v* in *D* as pa_*D*_(*v*). We write *e*_*ij*_(*D*) = **1***{*(*i, j*) ∈ *E*(*D*)*}* for the indicator that *D* has a directed edge from node *i* to node *j*.

We consider a set of random variables **Y** = *{Y*_1_, *Y*_2_, …, *Y*_*m*_*}*, each random variable corresponding to a node in *V* . We assume that the probability distribution of **Y**, *P* (**Y** = **y**), obeys the local Markov property described by *D*, implying the joint distribution of **Y** can be factorized as follows [30]:

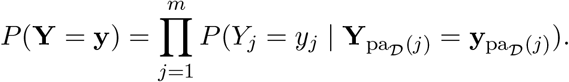

Following Choi *et al*. [13], we model each conditional distribution in this factorization as a negative binomial regression of *Y*_*j*_ on the counts of its parents, with

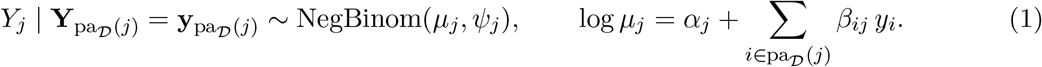

Here, NegBinom(*µ*_*j*_, *ψ*_*j*_) denotes a negative binomial distribution with mean *µ*_*j*_ and overdispersion *ψ*_*j*_, *α*_*j*_ an intercept term, and *β*_*ij*_ the direct effect of node *i* on the log mean of node *j*. Specifically,

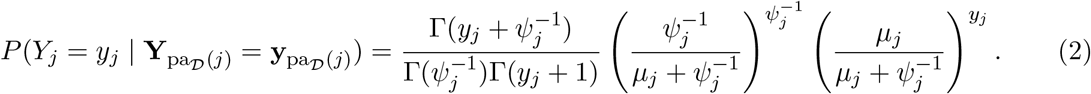

Under this parameterization, 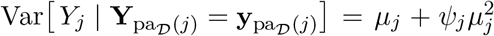 . We additionally let Θ = *{β*_*ij*_ : *i, j* ∈ *V, i* ≠ *j}* ∪ *{α*_*j*_, *ψ*_*j*_ : *j* ∈ *V }* collect the regression parameters.

Now consider a dataset of *N* observations, indexed *n* = 1, …, *N* . Let 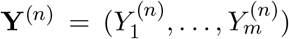denote the counts at the *m* nodes in observation *n*, **y**^(*n*)^ its realization, and *Y* = *{***y**^(1)^, …, **y**^(*N*)^*}* the full dataset. We assume our observations are independent, so the likelihood factorizes as

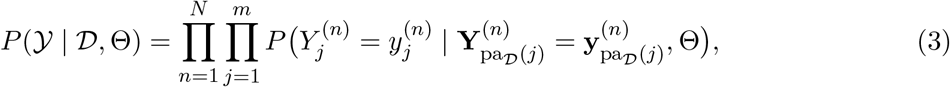

with each factor given by (2). In (3) we write the dependence on *D* and Θ explicitly, because we assign priors to each parameter in Θ below and a graph prior to *D* in Section 4.3.

We next place a prior on each of the regression parameters in Θ. As in Choi *et al*. [13], we let

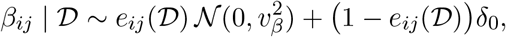

independently across ordered pairs *i* ≠*j*, where δ_0_ denotes a point mass at 0. Thus, the direct effect *β*_*ij*_ is drawn from a normal prior when the edge (*i, j*) is present in *D*, and fixed at 0 otherwise.

For the intercepts *α*_*j*_, we let

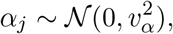

independently across *j* ∈ *V* . Finally, for the overdispersion parameters *ψ*_*j*_, we let

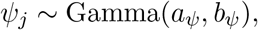

independently across *j* ∈ *V* .

The joint prior on Θ given *D* therefore factorizes as

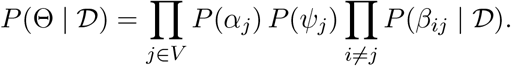

### 4.2 Topological prior

Each ordered pair of distinct nodes in *{*(*i, j*) ∈ *V × V* : *i* ≠ *j}* may be an edge in *D*. Suppose that prior evidence from one of *K* sources supports some of these pairs as edges. Furthermore, suppose some pairs can be ruled out as edges altogether. In the topological prior *T*, we assign each pair a single label, indicating whether the pair is forbidden, unsupported, or supported by a particular source of evidence.

Formally, the topological prior is the map

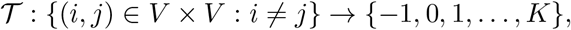

which assigns each ordered pair of distinct nodes (*i, j*) a label *T*_*ij*_ = *T* ((*i, j*)). A label *T*_*ij*_ = −1 marks the pair as forbidden, so (*i, j*) is excluded from being an edge in *D*. A label *T*_*ij*_ = 0 places (*i, j*) in the no-evidence class, meaning no source supports it. A label *T*_*ij*_ = *k* ∈ *{*1, …, *K}* places (*i, j*) in the *k*th evidence class, meaning (*i, j*) is supported by the *k*th evidence source. While the classes *k* = 0, 1, …, *K* are interchangeable in this model, we use 0 to indicate the no-evidence class by convention. We treat *T* as a fixed and known input throughout, constructed before inference independently from the data.

*T* partitions all ordered pairs of distinct nodes into the classes

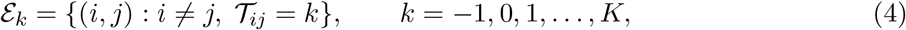

where *E*_−1_ collects the forbidden pairs and *E*_0_, …, *E*_*K*_ partition the remaining pairs, which we call permitted.

### 4.3 Graph prior

We give each class of the partition (4) its own edge probability. For *k* = 0, 1, …, *K*, let *ρ*_*k*_ be the edge probability of class *k*, and set *ρ*_−1_ = 0 for the forbidden class. We let ***ρ*** = (*ρ*_0_, *ρ*_1_, …, *ρ*_*K*_) collect the class edge probabilities of the *K* + 1 permitted classes.

Our graph prior includes each ordered pair *i* ≠ *j* as a directed edge independently with probability *ρ*_*Tij*_, and keeps the resulting graph only if it is acyclic,

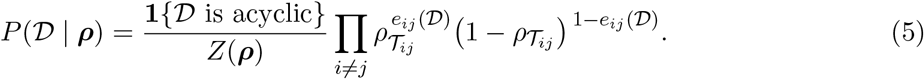

The normalizing constant

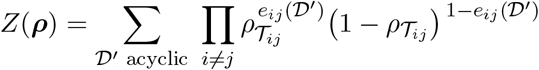

sums over all DAGs *D*^*′*^ on *V* . It is the probability that the graph is acyclic when each ordered pair *i* ≠*j* is included as a directed edge independently with probability *ρ*_*Tij*_ .

The product in (5) takes one factor per ordered pair, but it depends on *D* only through the number of edges present in each class. Writing

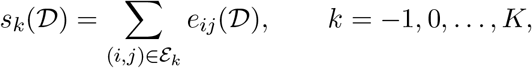

for the number of pairs in class *k* present as edges in *D*, and writing |*E*_*k*_| for the total number of pairs in the class, we may group the factors of (5) by class to obtain

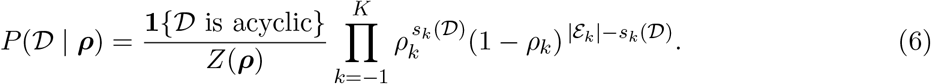

Notice because *ρ*_−1_ = 0, *P* (*D* | ***ρ***) = 0 if *s*_−1_(*D*) ≥ 1. The graph prior is thus positive only when no forbidden pairs are included as edges.

We place the prior

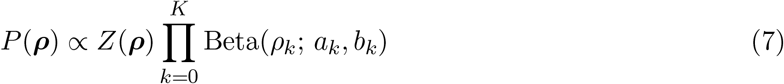

on the edge probabilities of the permitted classes, where Beta(*x*; *a, b*) denotes the density of the Beta(*a, b*) distribution evaluated at *x*, and *a*_*k*_, *b*_*k*_ *>* 0. The Beta densities express our prior belief about how likely a pair in each class is to be an edge, before we require the graph to be acyclic.

*Z*(***ρ***) cancels in the product *P* (*D* | ***ρ***) *P* (***ρ***). Similar approaches for canceling out intractable normalizing constants have been explored in previous work [13, 45]. The class edge probabilities given *D* therefore follow independent Beta distributions,

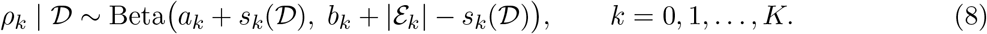

### 4.4 Posterior distribution

Our posterior distribution is defined over the DAG *D*, the regression parameters Θ, and the class edge probabilities ***ρ***. Given the observed counts *Y*,

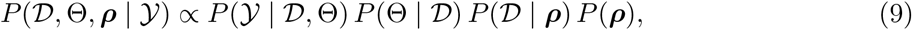

where each factor is specified in the preceding sections.

### 4.5 Markov chain Monte Carlo

We sample from the joint posterior in (9) using a parallel-tempering MCMC sampler modified from that of Choi *et al*. [13]. We run several MCMC chains in parallel, ordered along a ladder of temperatures. Higher temperature chains flatten the posterior distributions they target, making it easier to explore the parameter space, while the coldest chain targets (9) itself. At each iteration, the sampler attempts a replica-exchange swap between one adjacent pair of chains on the temperature ladder with a fixed swap probability. Otherwise, every chain applies, in turn, a Metropolis-Hastings move that adds, deletes, or reverses a single edge of *D*; a node-by-node Metropolis sweep over the regression parameters Θ; and a Gibbs draw of the class edge probabilities ***ρ*** from (8). All posterior inference uses post-burn-in draws from the cold chain. The MCMC algorithm is detailed in Supplementary Note 1.

### 4.6 Posterior inference

Let 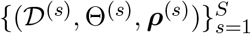be the *S* draws retained from the cold chain after burn-in. We define the posterior inclusion probability (PIP) of each ordered pair (*i, j*) as

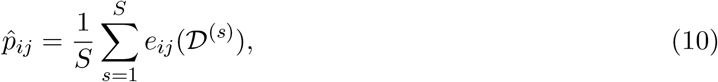

the proportion of draws whose DAG contains the edge (*i, j*). We summarize each direct effect *β*_*ij*_ by its posterior median and 95% central credible interval over the draws that contain the edge (*i, j*). We summarize each node intercept *α*_*j*_, overdispersion *ψ*_*j*_, and class edge probability *ρ*_*k*_ by its posterior median and 95% central credible interval.

### 4.7 Monocyte isolation, stimulation, and 10x Multiome profiling

CD14^+^ monocytes were isolated from healthy donor blood (Research Donors Ltd., ethical approval: NHS Research Ethics Committee ref. 25/LO/0213) by Ficoll density gradient centrifugation followed by fluorescence-activated cell sorting (FACS). Cells were cultured with macrophage colony-stimulating factor (M-CSF) for 16h and submitted to the 10x Multiome assay. Further details are provided in Supplementary Note 2.

### 4.8 10x Multiome data processing

#### snRNA-seq processing

We removed ambient RNA with CellBender v0.3.0 [46], using default settings except for the expected number of cells, which was set to the estimate from the Cell Ranger ARC pipeline. CellBender assigns each barcode a posterior cell probability, and we kept barcodes whose cell probability was above 0.5. Using Scanpy [47], we kept cells with at least 100 and fewer than 4,000 detected genes (as cells with more detected genes are likely doublets). We then removed genes detected in fewer than three cells and finally filtered out cells with 30% or more of their reads resulting from mitochondrial genes. We further identified and removed doublets among the remaining cells using SOLO [48] from scVI-tools. We then scaled the counts of each cell to sum to 10,000 using Scanpy and log-transformed them. We removed mitochondrial and ribosomal genes.

#### Defining metacells

To mitigate the sparse and noisy nature of 10x Multiome measurements, we combined the counts of cells with similar transcriptional profiles into metacells using SEACells [49] based on the similarity of the top 40 principal components of their normalized RNA-seq counts across cells. We set the number of metacells to approximately one per 38 cells, giving 83 metacells.

#### scATAC-seq processing

Fragment files from Cell Ranger ARC, with reads aligned to GRCh38, were processed using ArchR [50]. Cells with ≥ 1,000 unique fragments and transcription start site (TSS) enrichment score ≥ 1.2 were kept. scATAC-seq doublets were removed using ArchR’s built-in tool. ATAC-seq data for cells whose respective snRNA-seq profiles did not pass quality control (see “snRNA-seq processing”) were discarded. Chromatin accessibility was then quantified over a reference set of ATAC peaks derived from multiple 10x Multiome experiments (Supplementary Data 1). We restricted this peak set to peaks overlapping an ENCODE-annotated candidate enhancer (a proximal or distal enhancer-like signature, pELS or dELS) [51] and refer to these as enhancer peaks.

#### Building the metacell count matrix

The raw scATAC-seq counts per cell were grouped into a metacell by enhancer peak count matrix (see “Defining metacells”). Together, this yields a metacell by feature count matrix spanning genes (scRNA-seq) and enhancer peaks (scATAC-seq), comprising 83 metacells, which we refer to as the M-CSF 16h dataset. We refer to each gene or enhancer peak as a feature, and from here on to enhancer peaks simply as enhancers.

### 4.9 Selecting relevant features from the M-CSF 16h dataset

We selected a subset of gene and enhancer features from the M-CSF 16h dataset to use for SPARK network inference. We first kept all features from this dataset detected in more than 10% of metacells. Among these, we obtained the top 1,000 genes based on the highly variable gene score computed with Seurat’s `FindVariableFeatures`; under the variance-stabilizing transformation (`vst`;) [52], and refer to these as highly variable genes. We obtained the top 5,000 enhancers based on the sum of their counts across cells, and refer to these as highly accessible enhancers. We selected all features from these two sets.

We centered feature selection on “immGWAS” enhancers, whose overlapping pELS or dELS element contains a fine-mapped variant from an immune disease GWAS credible set (Supplementary Note 3) in the Open Targets Platform [15]. We added all immGWAS enhancers among the highly accessible enhancers to our feature set, along with three further categories of features. Table 2 lists each category and the number of features it contributes. First, we selected all highly variable genes whose transcription start sites (TSSs) lie within 1 Mb of the midpoint of an immGWAS enhancer, which we refer to as enhancer-proximal genes. Second, we selected all highly variable genes encoding transcription factors (TFs), as annotated by the Gene Ontology term `GO:0003700`; or any of its descendant terms [53, 54]. Finally, we selected all highly accessible enhancers whose midpoints lie within 1 Mb of the TSS of an enhancer-proximal gene or a TF, which we refer to as upstream enhancers. In total, we selected 377 features (72 genes and 305 enhancers).

**Table 2.** Features selected from the M-CSF 16h dataset.

| Feature type | Criterion for inclusion | Number | Overlap with features already selected |
| --- | --- | --- | --- |
| immGWAS enhancers | Highly accessible enhancers whose overlapping pELS or dELS element contains a fine-mapped variant from an immune disease GWAS credible set | 14 |  |
| Enhancer-proximal genes | Highly variable genes whose TSS lies within 1 Mb of the midpoint of an immGWAS enhancer | 17 |  |
| TFs | Highly variable genes annotated with `GO:0003700` or any of its descendant terms | 57 | 2 enhancer-proximal genes |
| Upstream enhancers | Highly accessible enhancers whose midpoint lies within 1 Mb of the TSS of an enhancer-proximal gene or a TF | 298 | 7 immGWAS enhancers |
| Genes |  | 72 |  |
| Enhancers |  | 305 |  |
| All features |  | 377 |  |

We then depth-normalized the counts of the selected features, separately for the genes and the enhancers, converting the counts of each metacell to proportions within that metacell and rescaling them to the median library size.

### 4.10 Constructing the topological prior

We constructed the topological prior *T* ^M-CSF^, which assigns each ordered pair in *{*(*i, j*) : *i* ≠ *j}* across the 377 selected features a label indicating which evidence source, if any, supports it. We used *K* = 4 evidence sources, two for TF to enhancer pairs and two for enhancer to gene pairs. Table 3 lists the number of pairs each source supports.

**Table 3.** Labels of the topological prior *T*^M-CSF^. Each pair supported by two sources received the label of the source we considered to be stronger evidence for a regulatory effect, so the labeled pairs equal the supported pairs minus the overlap.

| Label | Evidence source | Criterion | Supported<br>pairs | Overlap | Labeled<br>pairs |
| --- | --- | --- | --- | --- | --- |
| 1 | ChIP-Atlas | Binding site for the TF in the enhancer, with MACS2 $q < 10^{-5}$ | 207 | | 207 |
| 2 | JASPAR2024 CORE | Match to the TF’s motif in the enhancer, with $p < 5 \times 10^{-5}$ | 1,809 | 62 with ChIP-Atlas | 1,747 |
| 3 | Linear proximity | Enhancer overlapping the gene body, from its TSS to its transcription end site | 17 |  | 17 |
| 4 | Promoter Capture Hi-C | Contact in monocytes or macrophages between the enhancer and the gene’s promoter, with a CHiCAGO score of at least 5 | 55 | 4 with linear proximity | 51 |
| 0 | None | Permitted pair without support from any source |  |  | 15,785 |
| -1 | None | Gene to gene, enhancer to enhancer, outgoing from a gene that is not a TF, or enhancer to gene with the enhancer midpoint more than 1 Mb from the gene’s TSS |  |  | 123,945 |
| Total |  |  |  |  | 141,752 |

The first TF to enhancer source is TF binding from ChIP-Atlas [16]. We took ChIP-seq peaks in the antigen class ‘TFs and others’ with a MACS2 [55] significance of *q <* 10^−5^, from myeloid cells (CD14^+^ monocytes, 19 datasets; macrophages, 115 datasets) and myeloid cell lines (THP-1, 309 datasets; U-937, 50 datasets; Mono-Mac-6, 8 datasets; HL-60, 132 datasets). For each TF, we took the union of its peaks across all datasets. The second TF to enhancer source is motif matches from the JASPAR2024 CORE database [11]. We scanned enhancer peaks for motif matches with *p <* 5 *×* 10^−5^ using motifmatchr [56] via Signac [57], and mapped each motif to the genes encoding its TFs. We linked motifs for TF multimers (e.g., FOS::JUN) to each TF in the multimer.

The first enhancer to gene source is linear proximity. We linked an enhancer to a gene if the enhancer peak overlaps the gene body, from its TSS to its transcription end site, in the NCBI RefSeq annotation (GRCh38) [58]. The second enhancer to gene source is Promoter Capture Hi-C (PCHi-C) contacts in monocytes and macrophages (M0, M1, and M2) from Javierre et al. [12]. We lifted the published PCHi-C peak matrix over from GRCh37 to GRCh38 and kept contacts with a CHiCAGO [59] score of at least 5 in at least one of these cell types.

We considered a ChIP-seq peak, motif match, or PCHi-C contact to support an enhancer if it overlaps the enhancer peak. Pairs with support from two sources received the label of the source we considered to provide stronger evidence of a regulatory effect, ChIP-Atlas over JASPAR2024 and linear proximity over PCHi-C. We labeled with −1 every gene to gene and enhancer to enhancer pair, every outgoing pair from a gene that is not a TF, and every enhancer to gene pair whose enhancer midpoint lies more than 1 Mb from the gene’s TSS, prohibiting these pairs from appearing as edges in the sampled DAGs. We labeled all remaining pairs with 0, placing them in the no-evidence class.

### 4.11 Running SPARK network inference on the M-CSF 16h dataset

We ran SPARK on the normalized 83 *×* 377 count matrix of the M-CSF 16h dataset using the topological prior *T* ^M-CSF^. Each selected feature corresponds to a node *j* of the DAG, and its normalized counts across the 83 metacells are the observations of *Y*_*j*_.

In the prior (7) on ***ρ***, we set *a*_0_ = *b*_0_ = 1 for the no-evidence class. For each prior evidence source *k* = 1, …, 4, we elicited *a*_*k*_ and *b*_*k*_ by asking a domain expert for the 10th and 90th percentiles of their belief about how likely a pair in evidence class *E*_*k*_ is a regulatory effect (Table S1). In the priors on the regression parameters, we set the prior standard deviation of the direct effects to *v*_*β*_ = 0.05, the prior standard deviation of the intercepts to *v*_*α*_ = 2, and the shape and rate of the Gamma prior on the overdispersions to *a*_*ψ*_ = 1 and *b*_*ψ*_ = 2. We started every chain of the parallel-tempering sampler from the same DAG *D* and regression parameters Θ, obtained by regressing each node on its permitted parents with an *L*_1_-penalized negative binomial regression [60] and removing edges to break cycles (Supplementary Note 4). We ran 20 chains for 5,250,000 iterations, discarded the first 3,000,000 as burn-in, and kept every 10th draw of the cold chain, giving *S* = 225,000 draws for posterior inference. The remaining sampler settings are given in Table S5.

### 4.12 Simulation study

To assess how accurately SPARK reconstructs DAGs, we simulated count data from known regulatory networks. We restricted *T* ^M-CSF^ to the *m* = 51 nodes on chromosome 2 (5 TFs, 5 downstream genes, and 41 enhancers), and refer to this as *T* ^chr2^. Only two of the four evidence classes contained pairs among these nodes: 21 TF to enhancer pairs supported by JASPAR2024 CORE and 13 enhancer to gene pairs supported by Promoter Capture Hi-C. We relabeled these as classes *k* = 1 and *k* = 2, respectively. In total, *T* ^chr2^ had 302 permitted pairs, the 34 pairs in these two classes and 268 in the no-evidence class. The remaining 2,248 of the 2,550 ordered pairs were forbidden.

We first describe how we draw a single ground-truth DAG from *T* ^chr2^. We included each ordered pair *i* ≠ *j* as an edge independently with probability 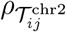, redrawing until the graph was acyclic. We used the values *ρ*_−1_ = 0, *ρ*_0_ = 0.05, *ρ*_1_ = 0.286, and *ρ*_2_ = 0.606, the last two being the means of the elicited Beta densities in Table S1.

We next describe how we simulate count data from a single ground-truth DAG. This is done based on SPARK’s likelihood, where the count of each node *j* follows a negative binomial distribution with overdispersion *ψ*_*j*_, whose log mean log *µ*_*j*_ is the intercept *α*_*j*_ plus the counts of its parents weighted by the direct effects *β*_*ij*_. Simulating counts therefore requires us to specify a value for each of these regression parameters. We set *ψ*_*j*_ = 0.1 for all *j* = 1, …, 51. We drew *α*_*j*_ ∼ *N* (log 2, 1) for all *j* = 1, …, 51 and *β*_*ij*_ ∼ *N* (0, 0.3^2^) for all ordered pairs *i* ≠ *j*, independently. We simulated each of the *N* = 50 observations separately, using the same procedure. The mean count of a node depends on the counts of its parents, so for each observation, we drew the node counts one at a time in an order where every node comes after its parental nodes in the DAG. Nodes without parents were drawn first from a negative binomial distribution with mean exp(*α*_*j*_) and overdispersion *ψ*_*j*_ = 0.1, and every later node was drawn once the counts of all its parents were available. Although we drew *β*_*ij*_ for every ordered pair, only the *β*_*ij*_ corresponding to the edges present in the DAG were used during this procedure.

We repeated these two steps, drawing a ground-truth DAG and simulating count data from it, until we had ten ground-truth DAGs, each with a simulated dataset. During this procedure, count simulation for one DAG produced a mean count *µ*_*j*_ above 10^7^ for at least one node and observation. We therefore discarded this DAG and its dataset, and replaced them with a newly drawn DAG and dataset whose mean count *µ*_*j*_ did not exceed 10^7^ for any node and observation. We drew *α*_*j*_ and *β*_*ij*_ only once, before drawing any DAG, and used them for all ten DAGs, so a pair that is an edge in several DAGs has the same direct effect in each.

We ran SPARK on each of the ten simulated datasets under four prior settings, which we call (1) non-informative, (2) expert-elicited, (3) strongly informative but misspecified, and (4) single class. In settings (1) to (3), SPARK was given *T* ^chr2^, the same topological prior from which the ground-truth DAGs were drawn. In setting (4), SPARK was given *T* ^chr2^ with the pairs of both evidence classes relabeled 0, so all 302 permitted pairs belonged to the no-evidence class, while the forbidden pairs remained forbidden.

The hyperparameters of the prior on ***ρ*** were as follows. In all four settings, we set *a*_0_ = *b*_0_ = 1 for the no-evidence class. For setting (1), we set *a*_*k*_ = *b*_*k*_ = 1 for *k* = 1, 2. For setting (2), we set (*a*_1_, *b*_1_) = (2.20, 5.50) and (*a*_2_, *b*_2_) = (5.68, 3.70), the elicited values for JASPAR2024 CORE and Promoter Capture Hi-C in Table S1, giving Beta densities with means of 0.286 and 0.606, equal to the generating values. For setting (3), we set (*a*_1_, *b*_1_) = (16.8, 4.2) and (*a*_2_, *b*_2_) = (3.25, 9.75), giving means of 0.80 and 0.25, far from the generating values. These Beta densities are narrow, with central 95% intervals of [0.61, 0.94] for *ρ*_1_ and [0.07, 0.51] for *ρ*_2_, neither of which contains the generating value. Setting (4) has only the no-evidence class, so only *a*_0_ and *b*_0_ are needed. In the priors on the regression parameters, we set the prior standard deviation of the direct effects to *v*_*β*_ = 0.2, the prior standard deviation of the intercepts to *v*_*α*_ = 2, and the shape and rate of the Gamma prior on the overdispersions to *a*_*ψ*_ = 1 and *b*_*ψ*_ = 2. We started every chain of the parallel-tempering sampler from the empty graph, ran 10 chains for 250,000 iterations, discarded the first 150,000 as burn-in, and kept every draw of the cold chain, giving *S* = 100,000 draws for posterior inference. The remaining sampler settings are given in Table S6.

### 4.13 Benchmarking against other methods

We compared SPARK’s edge recovery on the ten simulated datasets with that of two other methods, DAG0 [13] and the Bayesian ridge regression in CellOracle’s network inference step [14].

We ran DAG0 using the function `mcmc 1sampleDAG0`; from the BayesDAG0 R package. We started the DAG0 sampler from the empty graph and set its number of chains, temperature ladder, swap probability, number of iterations, burn-in, and prior on the overdispersions *ψ*_*j*_ identically to the SPARK runs. We took its proposal step sizes and remaining priors from the package’s worked example. We calculated the PIP of each ordered pair as the proportion of retained draws from DAG0’s cold chain whose DAG contains the edge, as defined for SPARK. CellOracle reconstructs a network by fitting, for each gene, a regularized linear regression of its expression on the expression of the TFs that its base GRN permits as regulators. We reimplemented this regression with *T* ^chr2^ in place of the base GRN, so the permitted pairs are those of SPARK.

Unlike CellOracle’s networks, SPARK’s include enhancers as nodes, so we fit one regression per gene and per enhancer, with enhancers permitted as regulators of genes. We applied the transformation log_2_(1 + *y*) to the simulated counts, matching CellOracle’s use of log-transformed expression. For each node *j*, we regressed its log count on the log counts of the nodes *i* for which (*i, j*) is a permitted pair. Following CellOracle, we divided each predictor by its standard deviation across cells and left the response unscaled, fitting by Bayesian ridge regression with the `scikit-learn`; implementation and the default settings CellOracle uses. Writing 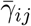and *σ*_*ij*_ for the posterior mean and standard deviation of the coefficient on node *i* in the regression for node *j*, and Φ(*·* ; *µ, σ*) for the cumulative distribution function of the normal distribution with mean *µ* and standard deviation *σ*, we scored each pair (*i, j*) by

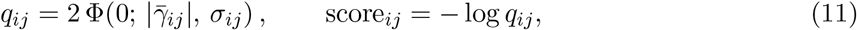

so that *q*_*ij*_ is twice the posterior probability that the coefficient takes the sign opposite to 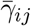. For SPARK, DAG0, and the regression, we ranked the 302 permitted pairs by their PIP or score

(11) and computed PR-AUC against each ground-truth DAG. All three methods were run on the same ten datasets, so their PR-AUC values are paired by ground-truth DAG.

## 5 Implementation

All analyses in this paper were carried out in Python, except DAG0, which we ran in R. The code that reproduces every analysis, figure, and table is available at https://github.com/nobuakimasaki/SPARK-eGRN-preprint.

## 6 Data and Resource Availability

The reference set of ATAC peaks (hg38) used to quantify chromatin accessibility (Supplementary Data 1), the metacell by feature count matrix of the M-CSF 16h dataset used as input to SPARK, the topological prior constructed for this dataset, the simulated datasets, and the outputs needed to regenerate every figure and table are available at https://github.com/nobuakimasaki/SPARK-eGRN-preprint.

## 7 Acknowledgments

We thank Ivan Andrew, Laurence Game (LMS Genomics Facility) and the members of LMS Bioinformatics Facility for 10x Multiome library preparation, sequencing, and initial processing. We thank S. Rong and all members of the Functional Gene Control Group for helpful suggestions and discussions. This work was supported by the British Heart Foundation project grant (PG/23/11350) and core funding from the Medical Research Council of the UK (MC-A652-5QA20) to M.S. L.B. gratefully acknowledges funding from the MRC (MR/W029790/1), the Alan Turing Institute under the Engineering and Physical Sciences Research Council (EPSRC) grant EP/N510129/1, the Marmaduke Sheild Fund, and the NIHR Cambridge Biomedical Research Centre (NIHR203312).

## Notes

https://github.com/nobuakimasaki/SPARK-eGRN-preprint

### Competing Interest Statement

The authors have declared no competing interest.

