## Supplementary Materials for "SPARK: Bayesian Structure Learning of Enhancer-based Gene Regulatory Networks from Single-cell Multiomics Data and Prior Knowledge"

**Supplementary Note 2. Details on monocyte isolation, stimulation, and 10x Multiome profiling** **7**

**Supplementary Note 3. Immune disease GWAS credible sets** **8**

**Supplementary Note 4. Warm-start initialization** **9**

**Supplementary Figures** **10**

**Supplementary Tables** **13**

**Supplementary References** **18**

### Supplementary Note 1. MCMC algorithm

We sample the DAG $\mathcal{D}$, the regression parameters $\Theta$, and the class edge probabilities $\boldsymbol{\rho}$ jointly from the posterior using a parallel-tempering MCMC sampler. Write $\boldsymbol{\xi} = (\mathcal{D}, \Theta, \boldsymbol{\rho})$ for the sampler state and $P(\boldsymbol{\xi}) = P(\mathcal{D}, \Theta, \boldsymbol{\rho} \mid \mathcal{Y})$ for the posterior evaluated at $\boldsymbol{\xi}$.

We run $R$ chains at temperatures $T_r = f^{\,r}$, $r = 0, \ldots, R-1$ using a geometric spacing factor $f > 1$, so that $T_0 = 1$. Chain $r$ targets the tempered posterior

$$\pi_r(\boldsymbol{\xi}) \propto P(\boldsymbol{\xi})^{1/T_r}, \tag{1}$$

so chain 0 targets $P(\boldsymbol{\xi})$ itself. Each chain $r$ holds its own state $\boldsymbol{\xi}_r$ and its untempered log-posterior $\ell_r = \log P(\boldsymbol{\xi}_r)$.

With probability $p_{\text{ex}}$ the sampler makes a swap move, exchanging the states of two chains with adjacent temperatures. Otherwise every chain applies, in order,

1. a graph update, proposing the birth, death, or reversal of a single edge of $\mathcal{D}$ and the direct effect $\beta_{ij}$ of any proposed edge;
2. a regression parameter update, a Metropolis step applied successively to each node's regression parameters;
3. an edge probability update, a Gibbs draw of the class edge probabilities from $P(\boldsymbol{\rho} \mid \mathcal{D})$.

We describe the swap move and these three updates in the subsections below, and write pseudocode for the full sampler in Algorithm 1 at the end of this section.

#### 1.1 Swap move

In the swap move, we propose exchanging the states of two chains adjacent on the temperature ladder, chains $r$ and $r+1$, so chain $r$ continues from state $\boldsymbol{\xi}_{r+1}$ and chain $r+1$ from state $\boldsymbol{\xi}_r$.

The proposal is accepted with probability

$$\min\Big\{1,\ \exp\Big[\Big(\frac{1}{T_r} - \frac{1}{T_{r+1}}\Big)\big(\log P(\boldsymbol{\xi}_{r+1}) - \log P(\boldsymbol{\xi}_r)\big)\Big]\Big\}, \tag{2}$$

which we compute from the log posteriors $\ell_r$ and $\ell_{r+1}$ stored by the two chains.

##### 1.2 Graph update

In the graph update, we apply a Metropolis-Hastings step that proposes the birth, death, or reversal of a single edge of $\mathcal{D}$, together with the direct effect of any edge it creates. Recall that $\beta_{ij}$, the direct effect of node $i$ on the log mean of node $j$, a priori follows $\mathcal{N}(0, v_\beta^2)$ when $(i,j)$ is an edge of $\mathcal{D}$ and equals 0 otherwise.

Write $\mathcal{P} = \bigcup_{k=0}^{K} \mathcal{E}_k$ for the ordered pairs the topological prior permits. Given the current DAG $\mathcal{D}$, each proposal type has its own set of candidate pairs,

$$\begin{aligned} \mathcal{C}_{\text{birth}}(\mathcal{D}) &= \big\{(i,j) \in \mathcal{P} \setminus E(\mathcal{D}) \;:\; E(\mathcal{D}) \cup \{(i,j)\} \text{ is acyclic}\big\}, \\ \mathcal{C}_{\text{death}}(\mathcal{D}) &= E(\mathcal{D}), \\ \mathcal{C}_{\text{reversal}}(\mathcal{D}) &= \big\{(i,j) \in E(\mathcal{D}) \;:\; (j,i) \in \mathcal{P},\ \big(E(\mathcal{D}) \setminus \{(i,j)\}\big) \cup \{(j,i)\} \text{ is acyclic}\big\}, \end{aligned} \tag{3}$$

of sizes $n_{\text{birth}}$, $n_{\text{death}}$, and $n_{\text{reversal}}$ respectively.

Let $\mathcal{M} = \big\{m \in \{\text{birth}, \text{death}, \text{reversal}\} : \mathcal{C}_m(\mathcal{D}) \neq \emptyset\big\}$ be the proposal types with at least one candidate. The proposal draws a type $m$ uniformly from $\mathcal{M}$, then a pair $(i,j)$ uniformly from $\mathcal{C}_m(\mathcal{D})$, and, for a birth or reversal, a direct effect $\tilde\beta \sim \mathcal{N}(0, v_\beta^2)$ for the edge it creates. Writing $\boldsymbol{\xi} = (\mathcal{D}, \Theta, \boldsymbol{\rho})$ for the current state, the proposed state $\boldsymbol{\xi}' = (\mathcal{D}', \Theta', \boldsymbol{\rho})$ is given by

$$\begin{aligned} \text{birth:}\quad & E(\mathcal{D}') = E(\mathcal{D}) \cup \{(i,j)\}, && \text{with } \beta_{ij} = \tilde\beta \text{ in } \Theta'; \\ \text{death:}\quad & E(\mathcal{D}') = E(\mathcal{D}) \setminus \{(i,j)\}, && \text{with } \beta_{ij} = 0 \text{ in } \Theta'; \\ \text{reversal:}\quad & E(\mathcal{D}') = \big(E(\mathcal{D}) \setminus \{(i,j)\}\big) \cup \{(j,i)\}, && \text{with } \beta_{ij} = 0 \text{ and } \beta_{ji} = \tilde\beta \text{ in } \Theta', \end{aligned}$$

all other elements of $\Theta$ unchanged. Writing $\phi(\cdot)$ for the $\mathcal{N}(0, v_\beta^2)$ density, the proposal distribution is

$$q(\boldsymbol{\xi}' \mid \boldsymbol{\xi}) = \frac{1}{|\mathcal{M}|} \begin{cases} \phi(\tilde\beta)/n_{\text{birth}} & \text{for a birth,} \\ 1/n_{\text{death}} & \text{for a death,} \\ \phi(\tilde\beta)/n_{\text{reversal}} & \text{for a reversal.} \end{cases} \tag{4}$$

The reverse of a birth is a death, of a death a birth, and of a reversal a reversal, so $q(\boldsymbol{\xi} \mid \boldsymbol{\xi}')$ is (4) evaluated on the proposed DAG $\mathcal{D}'$, with the direct effect of any restored edge taken to be the value $\beta_{ij}$ it held in $\mathcal{D}$.

The proposal is accepted with probability

$$\min\Big\{1,\ \exp\Big[\frac{1}{T_r}\big(\log P(\boldsymbol{\xi}') - \log P(\boldsymbol{\xi})\big) + \log q(\boldsymbol{\xi} \mid \boldsymbol{\xi}') - \log q(\boldsymbol{\xi}' \mid \boldsymbol{\xi})\Big]\Big\}. \tag{5}$$

Write $\Delta = \log P(\boldsymbol{\xi}') - \log P(\boldsymbol{\xi})$ for the untempered log-posterior difference in (5). The update leaves $\boldsymbol{\rho}$ unchanged, so this decomposes as $\Delta = \Delta_{\text{lik}} + \Delta_{\text{param}} + \Delta_{\text{graph}}$, the log differences in the first three factors of the posterior of the main paper, namely the likelihood $P(\mathcal{Y} \mid \mathcal{D}, \Theta)$, the parameter prior $P(\Theta \mid \mathcal{D})$, which the update changes only through the direct effects, and the graph prior $P(\mathcal{D} \mid \boldsymbol{\rho})$. Writing

$$L_j(\mathcal{D}, \Theta) = \sum_{n=1}^{N} \log P\big(Y_j^{(n)} = y_j^{(n)} \mid \mathbf{y}^{(n)}_{\mathrm{pa}_{\mathcal{D}}(j)}, \Theta\big) \tag{6}$$

for node $j$'s contribution to the log-likelihood, and abbreviating $L_j = L_j(\mathcal{D}, \Theta)$ and $L'_j = L_j(\mathcal{D}', \Theta')$, the three terms for each proposal type are

| | birth | death | reversal |
|---|---|---|---|
| $\Delta_{\text{lik}}$ | $L'_j - L_j$ | $L'_j - L_j$ | $(L'_j - L_j) + (L'_i - L_i)$ |
| $\Delta_{\text{param}}$ | $\log \phi(\tilde{\beta})$ | $-\log \phi(\beta_{ij})$ | $\log \phi(\tilde{\beta}) - \log \phi(\beta_{ij})$ |
| $\Delta_{\text{graph}}$ | $\operatorname{logit} \rho_{\mathcal{T}_{ij}}$ | $-\operatorname{logit} \rho_{\mathcal{T}_{ij}}$ | $\operatorname{logit} \rho_{\mathcal{T}_{ji}} - \operatorname{logit} \rho_{\mathcal{T}_{ij}}$ |

(7)

where $\beta_{ij}$ is the direct effect of the removed edge in $\mathcal{D}$. The last row holds because $\mathcal{D}$ and $\mathcal{D}'$ are both acyclic, so turning a pair $(i, j)$ from absent to present changes $\log P(\mathcal{D} \mid \boldsymbol{\rho})$ by $\log \rho_{\mathcal{T}_{ij}} - \log\big(1 - \rho_{\mathcal{T}_{ij}}\big) = \operatorname{logit} \rho_{\mathcal{T}_{ij}}$. Each factor of the posterior contributes only a few of its terms to (7), making $\Delta$ quicker to evaluate than $\log P(\boldsymbol{\xi}')$.

To form (3), we must check which permitted pairs can be added, and which existing edges can be reversed, without creating a cycle. We first form the reachability matrix $\mathbf{Z}$, with $Z_{ij} = 1$ when $\mathcal{D}$

contains a directed path from $i$ to $j$ and 0 otherwise. We fill $\mathbf{Z}$ one row at a time, row $i$ recording the descendants of node $i$, visited by a depth-first search from $i$ along the edges of $\mathcal{D}$. Writing $\mathbf{A}$ for the adjacency matrix of $\mathcal{D}$, whose entry $A_{ij}$ is 1 when $(i,j) \in E(\mathcal{D})$ and 0 otherwise, we also define a second matrix $\mathbf{Z}^{(2)}$ with entries

$$Z_{ij}^{(2)} = \bigvee_{l} \big(A_{il} \wedge Z_{lj}\big), \tag{8}$$

so $Z_{ij}^{(2)} = 1$ if and only if $\mathcal{D}$ contains a directed path from $i$ to $j$ of length at least two. Adding $(i,j)$ creates a cycle if and only if $Z_{ji} = 1$. Reversing $(i,j)$ creates a cycle when $i$ reaches $j$ by a path other than $(i,j)$, that is, by a path of length at least two, and so, if and only if $Z_{ij}^{(2)} = 1$.

##### 1.3 Regression parameter update

In the regression parameter update, we visit the nodes $j = 1, \ldots, m$, applying an independent Metropolis step to each. Write $\boldsymbol{\theta}_j = (\alpha_j, \boldsymbol{\beta}_{\cdot j}, \psi_j)$ for node $j$'s block of $\Theta$, where $\boldsymbol{\beta}_{\cdot j} = (\beta_{ij})_{i \in \mathrm{pa}_{\mathcal{D}}(j)}$ collects the direct effects on node $j$'s incoming edges. At node $j$ we draw a proposal $\boldsymbol{\theta}'_j$ from the symmetric Gaussian random walk

$$\alpha'_j = \alpha_j + \mathcal{N}(0, h_\alpha^2), \qquad \log \psi'_j = \log \psi_j + \mathcal{N}(0, h_\psi^2), \qquad \beta'_{ij} = \beta_{ij} + \mathcal{N}(0, h_\beta^2) \text{ for each } i \in \mathrm{pa}_{\mathcal{D}}(j), \tag{9}$$

drawn jointly, with fixed $h_\alpha, h_\beta, h_\psi$.

With $\mathcal{D}$ fixed, both the log-likelihood and the log-prior decompose over nodes,

$$\log P(\mathcal{Y} \mid \mathcal{D}, \Theta) = \sum_{j=1}^{m} L_j(\mathcal{D}, \Theta), \qquad \log P(\Theta \mid \mathcal{D}) = \sum_{j=1}^{m} \log P(\boldsymbol{\theta}_j \mid \mathcal{D}), \tag{10}$$

with $L_j$ the node log-likelihood (6) and

$$P(\boldsymbol{\theta}_j \mid \mathcal{D}) = P(\alpha_j)\ P(\psi_j) \prod_{i \in \mathrm{pa}_{\mathcal{D}}(j)} P(\beta_{ij} \mid \mathcal{D}) \tag{11}$$

node $j$'s block of the parameter prior. The acceptance probability is

$$\min\Big\{1,\ \exp\Big[\frac{1}{T_r}\big(L_j(\mathcal{D},\Theta') - L_j(\mathcal{D},\Theta) + \log P(\boldsymbol{\theta}'_j \mid \mathcal{D}) - \log P(\boldsymbol{\theta}_j \mid \mathcal{D})\big)\Big]\Big\}. \tag{12}$$

#### 1.4 Edge probability update

In the edge probability update, we draw the edge probability $\rho_k$ of each class, holding $\mathcal{D}$ and $\Theta$ fixed. From the graph prior and the prior on $\boldsymbol{\rho}$ (Section 4.3 of the main paper), we know that

$$\rho_k \mid \mathcal{D} \sim \mathrm{Beta}\big(a_k + s_k(\mathcal{D}),\ \ b_k + |\mathcal{E}_k| - s_k(\mathcal{D})\big), \qquad k = 0, 1, \ldots, K, \tag{13}$$

where $s_k(\mathcal{D})$ counts the pairs in class $k$ that are edges of $\mathcal{D}$, out of the $|\mathcal{E}_k|$ pairs in the class. Chain $r$ targets the tempered posterior (1), which raises this conditional to the power $1/T_r$,

$$\pi_r(\rho_k \mid \mathcal{D}) \propto \rho_k^{(a_k+s_k(\mathcal{D})-1)/T_r}\,(1-\rho_k)^{(b_k+|\mathcal{E}_k|-s_k(\mathcal{D})-1)/T_r}. \tag{14}$$

Therefore, under $\pi_r$,

$$\rho_k \mid \mathcal{D} \ \sim\ \mathrm{Beta}\Big(\big(a_k + s_k(\mathcal{D}) - 1\big)/T_r + 1,\ \ \big(b_k + |\mathcal{E}_k| - s_k(\mathcal{D}) - 1\big)/T_r + 1\Big), \qquad k = 0, 1, \ldots, K, \tag{15}$$

reducing to (13) when $T_r = 1$. We draw each $\rho_k$ independently from (15).

#### 1.5 MCMC sampler

Algorithm 1 collects the swap move and the three updates of Supplementary Notes 1.1 to 1.4 into the full parallel-tempering sampler, where **parfor** indicates a parallel for loop whereas **for** indicates a sequential for loop.

**Algorithm 1** Parallel-tempered MCMC for SPARK.

**Input:** data $\mathcal{Y}$, topological prior $\mathcal{T}$, hyperparameters $\left(v_\alpha, v_\beta, a_\psi, b_\psi, \{a_k, b_k\}_{k=0}^{K}\right)$, proposal step sizes $(h_\alpha, h_\beta, h_\psi)$, temperatures $1 = T_0 < \cdots < T_{R-1}$, swap probability $p_{\text{ex}}$, and number of iterations $n_{\text{iter}}$

Initialize the state $\boldsymbol{\xi}_r^{(0)} = \left(\mathcal{D}_r^{(0)}, \Theta_r^{(0)}, \boldsymbol{\rho}_r^{(0)}\right)$ of every chain $r = 0, \ldots, R-1$
**for** $t$ in $1, \ldots, n_{\text{iter}}$ **do**
  Draw a Bernoulli random variable $u$ with probability $p_{\text{ex}}$
  **if** $u = 1$ **then**
    Randomly select a pair of adjacent chains $(r, r+1)$ for $r \in \{0, \ldots, R-2\}$
    Perform a swap step to exchange $\boldsymbol{\xi}_r^{(t)}$ and $\boldsymbol{\xi}_{r+1}^{(t)}$
  **else**
    **parfor** $r$ in $0, \ldots, R-1$ **do**
      Perform a graph move for chain $r$ to update $\left(\mathcal{D}_r^{(t)}, \Theta_r^{(t)}\right)$
      Perform a Metropolis step at each node to update $\Theta_r^{(t)}$
      Perform a Gibbs step to update $\boldsymbol{\rho}_r^{(t)}$
    **end parfor**
  **end if**
**end for**

**Output:** MCMC samples from the cold chain, $\left\{\boldsymbol{\xi}^{(t)}\right\}_{t=1}^{n_{\text{iter}}} = \left\{\boldsymbol{\xi}_0^{(t)}\right\}_{t=1}^{n_{\text{iter}}}$

### Supplementary Note 2. Details on monocyte isolation, stimulation, and 10x Multiome profiling

#### Monocyte isolation and culture

Experiments used healthy donor blood from Research Donors Ltd., sourced through Cambridge Bioscience (ethical approval: NHS Research Ethics Committee ref. 25/LO/0213). Peripheral blood mononuclear cells (PBMCs) were isolated by Ficoll density gradient centrifugation. Monocytes were purified from PBMCs by fluorescence-activated cell sorting (FACS) using a lineage-exclusion and positive-selection antibody panel based on positive-selection markers HLA-DR (APC; BD Cat. no. 560744), CD14 (PE; BioLegend Cat. no. 301806), and CD16 (BV421; BioLegend Cat. no. 302038) and FITC-conjugated lineage-negative markers CD3 (BioLegend Cat. no. 300306), CD19 (BioLegend Cat. no. 302206), CD20 (BioLegend Cat. no. 302303), CD56 (BioLegend Cat. no. 304603), and CD66b (BioLegend Cat. no. 305103); TruStain FcX (BioLegend Cat. no. 422301) was included to block nonspecific Fc-receptor binding. Following isolation, monocytes were cultured in X-VIVO 15 Serum-free Hematopoietic Cell Medium (Lonza Cat. no. 02-060Q) in the presence of 100 ng/ml macrophage colony-stimulating factor (M-CSF; PeproTech Cat. no. 300-25) for 16h.

##### Nuclei isolation for 10x Multiome assay

Following stimulation, floating cells were collected directly, while adherent cells were detached using pre-warmed (37 °C) cell detachment buffer (ThermoFisher Cat. no. 13151014) and combined with the floating fraction. Detachment was confirmed by microscopy and cells were gently dissociated using wide-bore tips to minimize shear stress. Cells were pelleted and resuspended in PBS + 0.04% BSA, and up to 40,000 cells were transferred to a fresh tube for nuclei extraction. The cells were spun down and lysed in chilled lysis buffer (10 mM Tris-HCl [pH 7.4], 10 mM NaCl, 3 mM $MgCl_2$, 0.1% Tween-20, 0.1% NP-40, 0.01% Digitonin, 1% BSA, 1 mM DTT, and 1 U/µl RNase inhibitor [Sigma-Aldrich Cat. no. 3335399001]) for 30 s on ice, followed by addition, without mixing, of chilled wash buffer (10 mM Tris-HCl [pH 7.4], 10 mM NaCl, 3 mM $MgCl_2$, 1% BSA, 0.1% Tween-20, 1 mM DTT, and 1 U/µl RNase inhibitor) and centrifugation. The nuclei pellet was washed once more with chilled diluted nuclei buffer (made from the 20X stock supplied by 10x Genomics, supplemented with 1 mM DTT and 1 U/µl RNase inhibitor) under the same low-disruption conditions and resuspended in chilled diluted nuclei buffer. Nuclei concentration and integrity were assessed by Trypan blue staining, targeting a concentration of ∼1,600 nuclei/µl to achieve a recovery of 5,000 nuclei per sample for downstream 10x Genomics Multiome library preparation.

##### 10x Multiome library preparation, sequencing, and primary processing

Nuclei were prepared for 10x Multiome profiling using a standardized nuclei-isolation protocol. Gel bead-in-emulsion (GEM) generation, library preparation, and sequencing were performed by the LMS Genomics facility according to 10x Multiome protocol. Raw sequencing reads were processed using Cell Ranger ARC (v2.0.0) with default parameters by the LMS Bioinformatics facility.

#### Supplementary Note 3. Immune disease GWAS credible sets

We downloaded the credible set dataset of the Open Targets Platform [1], which collects fine-mapped variants from both genome-wide association studies (GWAS) and molecular quantitative trait locus (QTL) studies, and kept only the GWAS credible sets. We considered a credible set to be of an immune disease when one of its study's Experimental Factor Ontology terms [2] is, or descends from, immune system disorder (`MONDO:0005046`) or asthma (`MONDO:0004979`) [3], and descends from neither neoplasm (`MONDO:0005070`) nor tonsillitis (`MONDO:0001039`).

### Supplementary Note 4. Warm-start initialization

The initial state $\boldsymbol{\xi}^{(0)}$ for the real data analysis is a warm start fitted via penalized regression. For each node $j$, we regress its counts on the counts of all nodes the topological prior $\mathcal{T}$ permits as its parents, that is, all $i \neq j$ with $\mathcal{T}_{ij} \neq -1$, using an $L_1$-penalized negative binomial regression. We fit these regressions with `glm1path` from the `mvabund` R package [4], rounding each response to integer counts. We select the value of the penalty parameter $\lambda$ minimizing BIC across the 25 values `glm1path` fits by default, and collect into a directed graph $\mathcal{G}$ every pair $(i, j)$ for which candidate parent $i$ has a nonzero fitted coefficient in the regression for node $j$.

Each regression is fitted separately, so $\mathcal{G}$ may not be acyclic. We therefore break its cycles greedily. We score each edge of $\mathcal{G}$ by the absolute value of its fitted coefficient and add edges to an empty graph in decreasing order of score, skipping any edge whose addition would create a directed cycle. The resulting acyclic graph is the initial DAG $\mathcal{D}^{(0)}$, its edges keep their fitted coefficients as the initial $\beta_{ij}$, and the fitted intercepts and overdispersions give the initial $\alpha_j$ and $\psi_j$. Nodes with no permitted parents start with $\alpha_j = 0.5$ and $\psi_j = 0.5$. On the multiome data from M-CSF stimulated monocytes, $\mathcal{G}$ has 1,279 edges, of which 5 are dropped to break cycles, resulting in an initial DAG $\mathcal{D}^{(0)}$ of 1,274 edges.

### Supplementary Figures

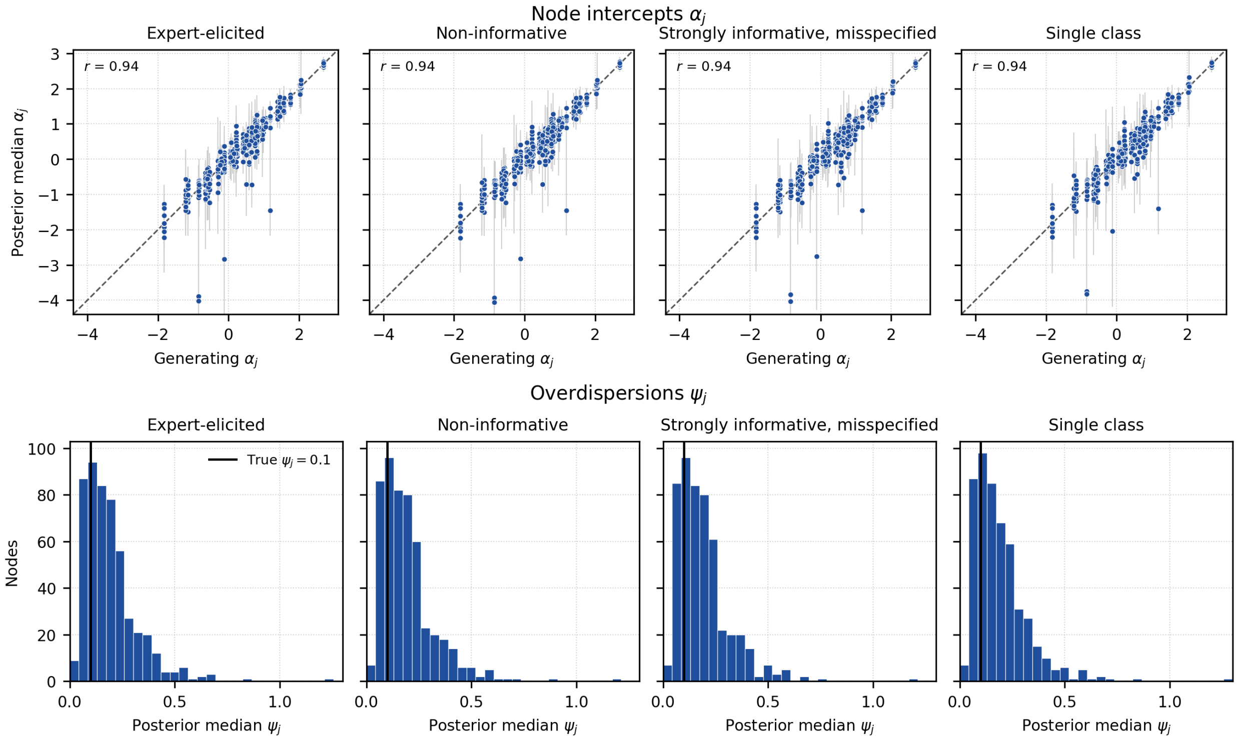

**Figure S1.** Recovery of the node intercepts and overdispersions on simulated data by SPARK under each of the four prior settings. The top row plots the posterior median of each node intercept $\alpha_j$ against its generating value, for 510 nodes pooled across DAGs, with its 95% central credible interval in gray. Each plot reports the Pearson correlation $r$ between the posterior medians and the generating values. The bottom row is a histogram of the posterior medians of the overdispersions $\psi_j$ over 510 nodes pooled across DAGs. The solid line marks the generating value $\psi_j = 0.1$, shared by every node.

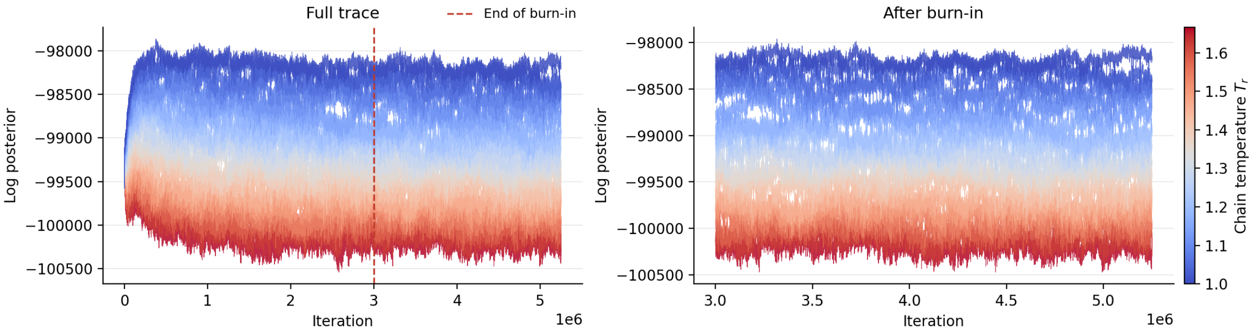

**Figure S2.** Log-posterior trace of each of the 20 chains in the real data analysis under the expert-elicited prior setting. The cold chain ($T_0 = 1$) targets the posterior itself, while hotter chains target flattened versions of it, which makes it easier to explore the space of DAGs. The left plot shows the full trace over all 5,250,000 iterations, with the end of the 3,000,000-iteration burn-in marked. The right plot shows the trace after burn-in.

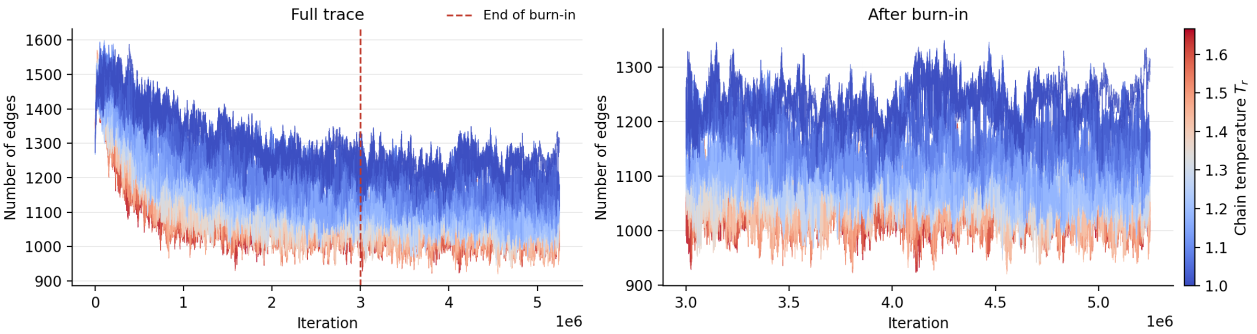

**Figure S3.** Trace of the number of edges in the sampled DAG for each of the 20 chains in the real data analysis under the expert-elicited prior setting. The cold chain ($T_0 = 1$) targets the posterior itself, while hotter chains target flattened versions of it, which makes it easier to explore the space of DAGs. The left plot shows the full trace over all 5,250,000 iterations, with the end of the 3,000,000-iteration burn-in marked. The right plot shows the trace after burn-in.

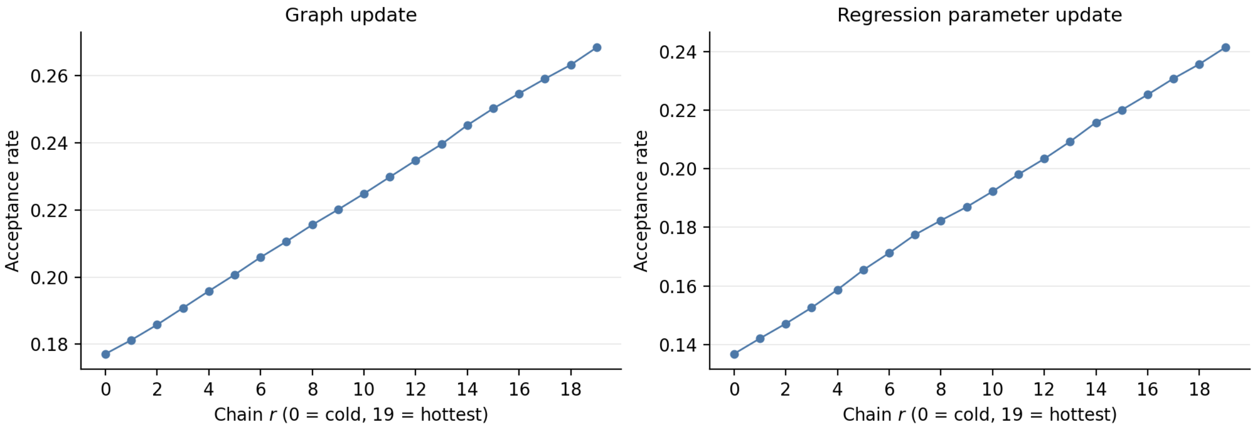

**Figure S4.** Acceptance rate of the graph update and the regression parameter update for each chain of the ladder, for the real data analysis under the expert-elicited setting. Rates are over the whole run, including burn-in.

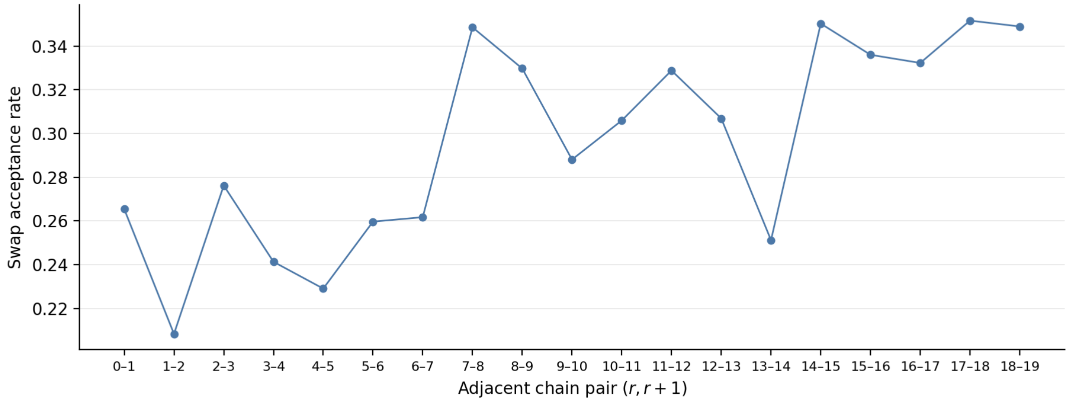

**Figure S5.** Swap acceptance rate for each adjacent pair of chains, for the real data analysis under the expert-elicited setting. Rates are over the whole run, including burn-in.

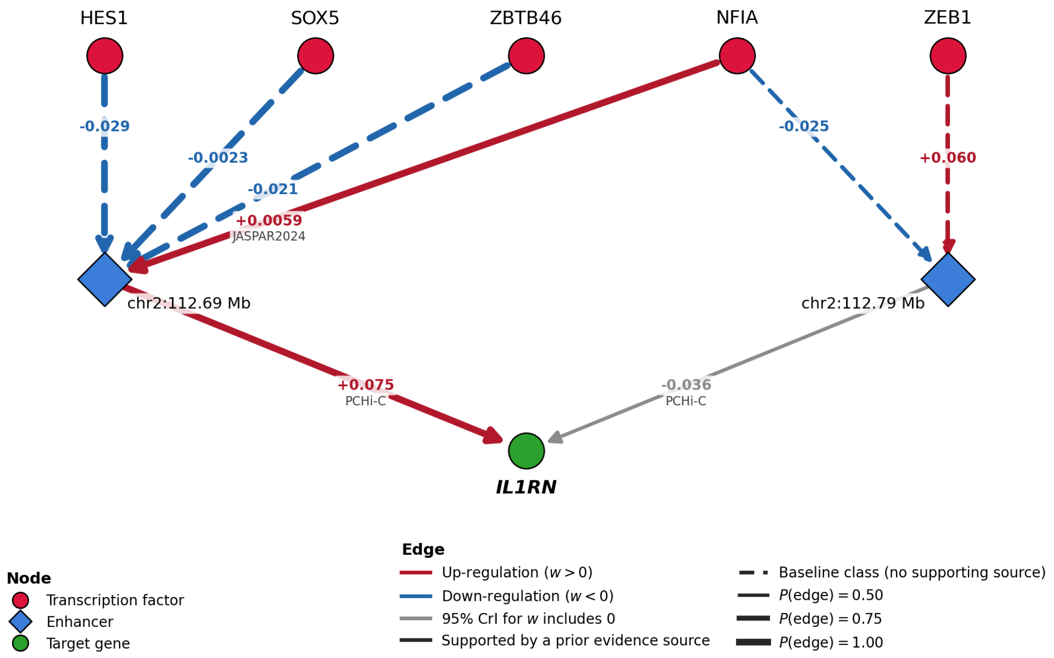

**Figure S6.** The cis-regulatory unit of *IL1RN* in the real data analysis under the expert-elicited prior setting, obtained by drawing every edge into *IL1RN* whose posterior inclusion probability (PIP) is at least 0.5, and then every such edge into the enhancers regulating *IL1RN*. Edge labels give the posterior median direct effect and, for prior-supported edges, the supporting evidence source. Edges are colored red or blue only when the 95% credible interval of the direct effect excludes 0.

### Supplementary Tables

| $k$ | Evidence source | $p_{10,k}$ | $p_{90,k}$ | $a_k$ | $b_k$ | Mean | Variance |
|---|---|---|---|---|---|---|---|
| 1 | ChIP-Atlas | 0.2 | 0.8 | 2.065 | 2.065 | 0.500 | 0.0487 |
| 2 | JASPAR2024 CORE | 0.1 | 0.5 | 2.204 | 5.505 | 0.286 | 0.0234 |
| 3 | TSS proximity | 0.5 | 0.9 | 5.505 | 2.204 | 0.714 | 0.0234 |
| 4 | Promoter Capture Hi-C | 0.4 | 0.8 | 5.680 | 3.700 | 0.606 | 0.0230 |

**Table S1.** Expert-elicited Beta densities in the prior on the class edge probabilities $\boldsymbol{\rho}$. We set $a_0 = b_0 = 1$ for the no-evidence class. For each prior evidence source $k = 1, \ldots, 4$, we elicited $a_k$ and $b_k$ by asking a domain expert (M.S.) for the 10th and 90th percentiles, $p_{10,k}$ and $p_{90,k}$, of their belief about how likely a pair in evidence class $\mathcal{E}_k$ is a regulatory effect. We chose $a_k$ and $b_k$ so that Beta$(a_k, b_k)$ has these percentiles. The last two columns are the mean and variance of Beta$(a_k, b_k)$.

| Method | DAG | True edges | Called | Correct | Precision | Recall | PR-AUC |
|---|---|---|---|---|---|---|---|
| SPARK, expert-elicited | 0 | 24 | 16 | 13 | 0.81 | 0.54 | 0.733 |
| | 1 | 22 | 14 | 12 | 0.86 | 0.55 | 0.733 |
| | 2 | 33 | 17 | 15 | 0.88 | 0.45 | 0.606 |
| | 3 | 20 | 12 | 11 | 0.92 | 0.55 | 0.701 |
| | 4 | 20 | 21 | 15 | 0.71 | 0.75 | 0.814 |
| | 5 | 29 | 20 | 16 | 0.80 | 0.55 | 0.717 |
| | 6 | 32 | 22 | 15 | 0.68 | 0.47 | 0.668 |
| | 7 | 23 | 14 | 12 | 0.86 | 0.52 | 0.751 |
| | 8 | 24 | 19 | 13 | 0.68 | 0.54 | 0.710 |
| | 9 | 25 | 16 | 13 | 0.81 | 0.52 | 0.726 |
| | Mean | 25.2 | 17.1 | 13.5 | 0.80 | 0.54 | 0.716 |
| SPARK, non-informative | 0 | 24 | 16 | 14 | 0.88 | 0.58 | 0.734 |
| | 1 | 22 | 16 | 14 | 0.88 | 0.64 | 0.722 |
| | 2 | 33 | 23 | 12 | 0.52 | 0.36 | 0.551 |
| | 3 | 20 | 16 | 11 | 0.69 | 0.55 | 0.702 |
| | 4 | 20 | 30 | 16 | 0.53 | 0.80 | 0.816 |
| | 5 | 29 | 21 | 17 | 0.81 | 0.59 | 0.719 |
| | 6 | 32 | 30 | 18 | 0.60 | 0.56 | 0.669 |
| | 7 | 23 | 22 | 13 | 0.59 | 0.57 | 0.742 |
| | 8 | 24 | 31 | 15 | 0.48 | 0.62 | 0.683 |
| | 9 | 25 | 29 | 16 | 0.55 | 0.64 | 0.702 |
| | Mean | 25.2 | 23.4 | 14.6 | 0.65 | 0.59 | 0.704 |
| SPARK, strongly informative, misspecified | 0 | 24 | 27 | 15 | 0.56 | 0.62 | 0.700 |
| | 1 | 22 | 30 | 14 | 0.47 | 0.64 | 0.726 |
| | 2 | 33 | 24 | 11 | 0.46 | 0.33 | 0.535 |
| | 3 | 20 | 27 | 11 | 0.41 | 0.55 | 0.639 |
| | 4 | 20 | 30 | 13 | 0.43 | 0.65 | 0.715 |
| | 5 | 29 | 32 | 14 | 0.44 | 0.48 | 0.611 |
| | 6 | 32 | 28 | 17 | 0.61 | 0.53 | 0.662 |
| | 7 | 23 | 25 | 13 | 0.52 | 0.57 | 0.712 |
| | 8 | 24 | 26 | 13 | 0.50 | 0.54 | 0.666 |
| | 9 | 25 | 30 | 16 | 0.53 | 0.64 | 0.673 |
| | Mean | 25.2 | 27.9 | 13.7 | 0.49 | 0.56 | 0.664 |

*Continued on next page*

| Method | DAG | True edges | Called | Correct | Precision | Recall | PR-AUC |
|---|---|---|---|---|---|---|---|
| SPARK, single class | 0 | 24 | 12 | 12 | 1.00 | 0.50 | 0.623 |
| | 1 | 22 | 15 | 13 | 0.87 | 0.59 | 0.733 |
| | 2 | 33 | 8 | 8 | 1.00 | 0.24 | 0.501 |
| | 3 | 20 | 9 | 9 | 1.00 | 0.45 | 0.588 |
| | 4 | 20 | 10 | 10 | 1.00 | 0.50 | 0.712 |
| | 5 | 29 | 13 | 12 | 0.92 | 0.41 | 0.620 |
| | 6 | 32 | 16 | 13 | 0.81 | 0.41 | 0.565 |
| | 7 | 23 | 9 | 8 | 0.89 | 0.35 | 0.614 |
| | 8 | 24 | 10 | 10 | 1.00 | 0.42 | 0.580 |
| | 9 | 25 | 11 | 10 | 0.91 | 0.40 | 0.656 |
| | Mean | 25.2 | 11.3 | 10.5 | 0.94 | 0.43 | 0.619 |
| DAG0 | 0 | 24 | 9 | 9 | 1.00 | 0.38 | 0.606 |
| | 1 | 22 | 9 | 9 | 1.00 | 0.41 | 0.697 |
| | 2 | 33 | 5 | 5 | 1.00 | 0.15 | 0.499 |
| | 3 | 20 | 5 | 5 | 1.00 | 0.25 | 0.595 |
| | 4 | 20 | 6 | 6 | 1.00 | 0.30 | 0.709 |
| | 5 | 29 | 5 | 5 | 1.00 | 0.17 | 0.582 |
| | 6 | 32 | 7 | 6 | 0.86 | 0.19 | 0.550 |
| | 7 | 23 | 6 | 6 | 1.00 | 0.26 | 0.580 |
| | 8 | 24 | 7 | 7 | 1.00 | 0.29 | 0.576 |
| | 9 | 25 | 8 | 8 | 1.00 | 0.32 | 0.666 |
| | Mean | 25.2 | 6.7 | 6.6 | 0.99 | 0.27 | 0.606 |
| Regularized regression (CellOracle) | 0 | 24 | 9 | 8 | 0.89 | 0.33 | 0.519 |
| | 1 | 22 | 10 | 8 | 0.80 | 0.36 | 0.611 |
| | 2 | 33 | 7 | 5 | 0.71 | 0.15 | 0.404 |
| | 3 | 20 | 5 | 4 | 0.80 | 0.20 | 0.362 |
| | 4 | 20 | 4 | 3 | 0.75 | 0.15 | 0.574 |
| | 5 | 29 | 9 | 7 | 0.78 | 0.24 | 0.457 |
| | 6 | 32 | 11 | 8 | 0.73 | 0.25 | 0.474 |
| | 7 | 23 | 4 | 4 | 1.00 | 0.17 | 0.543 |
| | 8 | 24 | 8 | 7 | 0.88 | 0.29 | 0.471 |
| | 9 | 25 | 8 | 7 | 0.88 | 0.28 | 0.499 |
| | Mean | 25.2 | 7.5 | 6.1 | 0.82 | 0.24 | 0.492 |

**Table S2.** Edge detection on simulated data for each of the ten ground-truth DAGs, by SPARK under each of the four prior settings, DAG0, and the regularized regression from CellOracle. Called is the number of pairs with a posterior inclusion probability of at least 0.5, or with a p-value below 0.001 for the regularized regression, the default cutoff used in CellOracle. Correct is the number of called pairs that are edges of the ground-truth DAG.

| Setting | DAG | Posterior median class edge probability [95% CrI] | | |
|---|---|---|---|---|
| | | JASPAR2024 (true value 0.286) | PCHi-C (true value 0.606) | No evidence (true value 0.05) |
| Expert-elicited | 0 | 0.35 [0.14, 0.62] | 0.59 [0.33, 0.84] | 0.07 [0.02, 0.13] |
| | 1 | 0.30 [0.10, 0.58] | 0.53 [0.29, 0.78] | 0.08 [0.04, 0.15] |
| | 2 | 0.46 [0.22, 0.71] | 0.59 [0.31, 0.83] | 0.02 [0.00, 0.07] |
| | 3 | 0.22 [0.06, 0.49] | 0.70 [0.45, 0.90] | 0.03 [0.01, 0.07] |
| | 4 | 0.38 [0.14, 0.66] | 0.70 [0.44, 0.90] | 0.03 [0.01, 0.08] |
| | 5 | 0.30 [0.10, 0.58] | 0.74 [0.48, 0.92] | 0.05 [0.02, 0.11] |
| | 6 | 0.52 [0.27, 0.76] | 0.57 [0.30, 0.83] | 0.05 [0.02, 0.11] |
| | 7 | 0.42 [0.18, 0.69] | 0.56 [0.29, 0.82] | 0.04 [0.01, 0.09] |
| | 8 | 0.49 [0.24, 0.74] | 0.62 [0.36, 0.86] | 0.02 [0.00, 0.06] |
| | 9 | 0.44 [0.20, 0.69] | 0.54 [0.28, 0.79] | 0.06 [0.02, 0.13] |
| | RMSE | 0.14 | 0.07 | 0.02 |
| Non-informative | 0 | 0.43 [0.15, 0.77] | 0.52 [0.18, 0.90] | 0.07 [0.02, 0.13] |
| | 1 | 0.43 [0.10, 0.81] | 0.44 [0.14, 0.80] | 0.08 [0.04, 0.15] |
| | 2 | 0.73 [0.34, 0.97] | 0.47 [0.11, 0.90] | 0.03 [0.00, 0.07] |
| | 3 | 0.28 [0.04, 0.73] | 0.79 [0.43, 0.99] | 0.03 [0.01, 0.07] |
| | 4 | 0.60 [0.18, 0.97] | 0.80 [0.45, 0.99] | 0.03 [0.01, 0.08] |
| | 5 | 0.34 [0.09, 0.77] | 0.84 [0.38, 0.99] | 0.06 [0.02, 0.11] |
| | 6 | 0.77 [0.42, 0.99] | 0.54 [0.14, 0.95] | 0.05 [0.02, 0.11] |
| | 7 | 0.61 [0.23, 0.97] | 0.48 [0.14, 0.96] | 0.03 [0.01, 0.08] |
| | 8 | 0.73 [0.37, 0.98] | 0.66 [0.28, 0.97] | 0.02 [0.00, 0.06] |
| | 9 | 0.75 [0.34, 0.98] | 0.47 [0.13, 0.82] | 0.07 [0.02, 0.14] |
| | RMSE | 0.33 | 0.15 | 0.02 |
| Strongly informative, misspecified | 0 | 0.74 [0.55, 0.90] | 0.32 [0.13, 0.55] | 0.07 [0.03, 0.13] |
| | 1 | 0.74 [0.54, 0.90] | 0.30 [0.13, 0.52] | 0.08 [0.04, 0.15] |
| | 2 | 0.82 [0.63, 0.94] | 0.28 [0.11, 0.52] | 0.03 [0.00, 0.08] |
| | 3 | 0.74 [0.53, 0.91] | 0.43 [0.23, 0.65] | 0.03 [0.01, 0.07] |
| | 4 | 0.78 [0.60, 0.91] | 0.42 [0.21, 0.65] | 0.03 [0.01, 0.08] |
| | 5 | 0.77 [0.57, 0.92] | 0.41 [0.20, 0.64] | 0.05 [0.02, 0.10] |
| | 6 | 0.81 [0.62, 0.94] | 0.28 [0.11, 0.51] | 0.05 [0.02, 0.11] |
| | 7 | 0.79 [0.60, 0.93] | 0.30 [0.12, 0.55] | 0.03 [0.01, 0.08] |
| | 8 | 0.80 [0.62, 0.93] | 0.35 [0.17, 0.59] | 0.02 [0.00, 0.05] |
| | 9 | 0.82 [0.63, 0.94] | 0.29 [0.11, 0.53] | 0.06 [0.02, 0.13] |
| | RMSE | 0.50 | 0.27 | 0.02 |
| Single class | 0 | | | 0.12 [0.06, 0.18] |
| | 1 | | | 0.11 [0.06, 0.18] |
| | 2 | | | 0.09 [0.04, 0.15] |
| | 3 | | | 0.07 [0.04, 0.14] |
| | 4 | | | 0.09 [0.04, 0.15] |
| | 5 | | | 0.10 [0.05, 0.16] |
| | 6 | | | 0.12 [0.06, 0.19] |
| | 7 | | | 0.09 [0.05, 0.16] |
| | 8 | | | 0.09 [0.05, 0.15] |
| | 9 | | | 0.12 [0.06, 0.19] |
| | RMSE | | | 0.05 |

**Table S3.** Estimation of class edge probabilities on simulated data by SPARK under the four prior settings. Each entry is the posterior median of the class edge probability, with its 95% central credible interval in brackets. The true value of each class edge probability, used to generate the ground-truth DAGs, is given in parentheses below the name of its class. For each setting, RMSE is the root mean squared difference between the posterior median and the true value over the ten DAGs.

| TF | Edges | Negative | Supported | TF | Edges | Negative | Supported |
|---|---|---|---|---|---|---|---|
| AFF3 | 133 | 119 (89.5) | 0 (0.0) | E2F7 | 3 | 3 (100.0) | 1 (33.3) |
| NFIA | 119 | 94 (79.0) | 42 (35.3) | EGR2 | 3 | 0 (0.0) | 0 (0.0) |
| PPARG | 87 | 70 (80.5) | 14 (16.1) | PPARD | 3 | 1 (33.3) | 2 (66.7) |
| HES1 | 31 | 28 (90.3) | 7 (22.6) | TFEC | 3 | 1 (33.3) | 0 (0.0) |
| SOX5 | 16 | 11 (68.8) | 0 (0.0) | ZBTB46 | 3 | 2 (66.7) | 0 (0.0) |
| BACH2 | 12 | 8 (66.7) | 5 (41.7) | ZEB1 | 3 | 2 (66.7) | 2 (66.7) |
| ZNF273 | 11 | 11 (100.0) | 0 (0.0) | ZNF366 | 3 | 3 (100.0) | 0 (0.0) |
| BHLHE41 | 10 | 3 (30.0) | 0 (0.0) | EBF1 | 2 | 2 (100.0) | 0 (0.0) |
| ZBTB16 | 10 | 5 (50.0) | 0 (0.0) | ETV6 | 2 | 0 (0.0) | 0 (0.0) |
| ZBTB7C | 9 | 5 (55.6) | 3 (33.3) | KLF12 | 2 | 2 (100.0) | 2 (100.0) |
| MEF2A | 8 | 2 (25.0) | 5 (62.5) | LITAF | 2 | 1 (50.0) | 0 (0.0) |
| BCL11A | 6 | 3 (50.0) | 1 (16.7) | MEF2C | 2 | 0 (0.0) | 1 (50.0) |
| EGR1 | 6 | 6 (100.0) | 4 (66.7) | NPAS1 | 2 | 1 (50.0) | 0 (0.0) |
| GLIS3 | 6 | 1 (16.7) | 0 (0.0) | NR3C2 | 2 | 1 (50.0) | 0 (0.0) |
| HIVEP3 | 6 | 3 (50.0) | 0 (0.0) | PLAGL1 | 2 | 2 (100.0) | 0 (0.0) |
| ATF3 | 5 | 0 (0.0) | 0 (0.0) | RORB | 2 | 2 (100.0) | 0 (0.0) |
| ATOH8 | 5 | 3 (60.0) | 0 (0.0) | STAT4 | 2 | 1 (50.0) | 0 (0.0) |
| CREB5 | 5 | 2 (40.0) | 0 (0.0) | ZBTB25 | 2 | 0 (0.0) | 0 (0.0) |
| GTF2IRD1 | 4 | 2 (50.0) | 0 (0.0) | ZNF280C | 2 | 1 (50.0) | 0 (0.0) |
| MITF | 4 | 4 (100.0) | 0 (0.0) | BATF | 1 | 1 (100.0) | 1 (100.0) |
| REL | 4 | 2 (50.0) | 0 (0.0) | SMAD3 | 1 | 1 (100.0) | 0 (0.0) |
| RUNX2 | 4 | 2 (50.0) | 2 (50.0) | TCF4 | 1 | 1 (100.0) | 1 (100.0) |
| BHLHE40 | 3 | 2 (66.7) | 0 (0.0) | TGIF2 | 1 | 1 (100.0) | 1 (100.0) |
| All TFs | 553 | 415 (75.0) | 94 (17.0) | | | | |

**Table S4.** Signs of the inferred TF-enhancer edges in the real data analysis under the expert-elicited prior setting. Each row is one of the 46 TFs with at least one edge to an enhancer whose posterior inclusion probability (PIP) is at least 0.5, sorted by the number of outgoing edges. The negative column counts the edges whose posterior median direct effect is negative. The supported column counts the edges supported by a prior evidence source (ChIP-Atlas or JASPAR2024). Percentages of the TF's edges are in parentheses.

| Setting | Symbol | Value |
|---|---|---|
| Temperature ladder | $T_r$ | $(1/0.6)^{r/19}$, $r = 0, \ldots, 19$ |
| Swap probability | $p_{\text{ex}}$ | 0.2 |
| Intercept random-walk step | $h_\alpha$ | 0.1 |
| Weight random-walk step | $h_\beta$ | 0.03 |
| Log-dispersion random-walk step | $h_\psi$ | 0.1 |

**Table S5.** Sampler settings for the real data analysis. $h_\alpha$, $h_\beta$, and $h_\psi$ are the random-walk standard deviations of the regression parameter update.

| Setting | Symbol | Value |
|---|---|---|
| Temperature ladder | $T_r$ | $2.5^{r/9}$, $r = 0, \ldots, 9$ |
| Swap probability | $p_{\mathrm{ex}}$ | 0.2 |
| Intercept random-walk step | $h_\alpha$ | 0.25 |
| Weight random-walk step | $h_\beta$ | 0.075 |
| Log-dispersion random-walk step | $h_\psi$ | 0.25 |

**Table S6.** Sampler settings for the simulation study. $h_\alpha$, $h_\beta$, and $h_\psi$ are the random-walk standard deviations of the regression parameter update.
